# Androgen receptor activation stabilizes a hybrid epithelial/mesenchymal phenotype in presence of Notch-Jagged signaling

**DOI:** 10.1101/2025.11.09.687414

**Authors:** Souvik Guha, R Soundharya, Baishakhi Tikader, Mohit Kumar Jolly

## Abstract

Metastasis is a key cause of mortality in cancer. It is driven majorly by clusters of circulating tumor cells which are often comprised of cells in a hybrid epithelial/mesenchymal (E/M) phenotype. Hybrid E/M phenotype has also been shown to be more tumor-initiating and therapy-resistant, but how cells maintain the hybrid E/M phenotype(s) remains an active area of investigation. Here, we develop a mathematical model that couples the intracellular dynamics of key players of Epithelial-Mesenchymal Transition (EMT) with the Androgen Receptor (AR), and cell-cell communication through Notch-Delta-Jagged signaling, in the context of prostate cancer. Our simulations show that AR can stabilize a hybrid E/M phenotype predominantly in the presence of Notch-Jagged, but not Notch-Delta, signaling. Implementing this model on a multi-cellular lattice, we observed that AR can alter the fraction of cells exhibiting a hybrid E/M and a mesenchymal phenotype. Finally, through analysis of transcriptomic and patient survival data, we found that the co-expression of AR and Jagged correlated with worse outcomes. Together, these results highlight the outcome of emergent dynamics between the Notch and AR signaling axis in promoting prostate cancer aggressiveness.

## 1 Introduction

Metastasis accounts for over 90% of cancer-associated mortality [1]. A key aspect of metastasizing cells is phenotypic plasticity which enables them to adapt to their dynamic microenvironments, while also evading different therapeutic interventions and attacks by the immune system [2]. Epithelial-to-mesenchymal transition (EMT) and its reverse – mesenchymal-to-epithelial transition – is a crucial axis of phenotypic plasticity that are also pivotal during embryonic development and wound healing[2]. EMT has also been linked to immune-suppressive and immune-evasive phenotypes through secreting cytokines such as IL-6 and IL-8 [3], and upregulation of checkpoint molecules such as PD-L1[4, 5]. The processes of EMT and MET are rarely binary; instead, cells can display mixed phenotypes with both epithelial and mesenchymal traits, enabling them capable of exhibiting collective cell migration[6]. Such collective migration can lead to the emergence of clusters of circulating tumor cells (CTCs) – the key drivers of metastasis. Clusters of CTC are rare and often contain only 5-8 cells, but can have up to 50 times more metastatic propensity, demonstrating their disproportionately high contribution to metastasis[7]. CTC clusters associate with worse clinical outcomes compared to individual CTCs that can be attributed to their relatively higher tumorigenic potential and/or reduced cell death in circulation [8]. Thus, decoding the intracellular and intercellular processes that enable the formation of CTC clusters is critical to eventually curb metastasis.

Broadly speaking, two fundamental conditions should be met for formation of clusters of CTCs. First, tumor cells forming CTC clusters should have the traits of both cell-cell adhesion and migration, which are usually exhibited by hybrid epithelial/mesenchymal (E/M) phenotypes[9, 10]. Second, these cells should be spatially co-localized in a primary tumor. One way to achieve this co-localization can be induction of hybrid E/M phenotypes in neighboring tumor cells and/or stabilization of hybrid E/M phenotypes by cell-cell communication. Our previous work has highlighted the role of Notch-Jagged signaling based cell-cell communication (juxtacrine) in stabilizing the hybrid E/M phenotypes[10]. Consistent with our modeling predictions, in breast cancer, the expression of Jagged was observed to be higher in CTC clusters, and its knockdown in SUM149 cells led to reduced organoid formation[11, 12]. However, how the crosstalk of Notch-Jagged signaling with additional context-specific molecules in other cancer types impacts this observation remains unclear.

Hybrid E/M phenotypes and CTC clusters have also been reported in prostate cancer (PCa)[13, 14]. PCa is fundamentally driven by androgen receptor (AR) signaling, thus androgen deprivation therapy (ADT) is widely used in its treatment. Although PCa patients initially respond to ADT and show clinical regression, most tumors can evolve mechanisms of resistance (including, but not limited to, mutated or spliced forms of AR such as AR-V7) and progress to castration-resistant prostate cancer (CRPC)[15]. Notch signaling is known to be activated in cell lines resistant to enzalutamide (AR antagonist) compared to the sensitive (parental) ones, and its abrogation in vitro and in vivo could restore the sensitivity of PCa cells to enzalutamide[16]. Consistently, inhibiting Notch in vitro via CRISPR or pharmacologically could reverse enzalutamide resistance[17]. Further, immunohistochemical analysis of PCa tumor samples revealed an upregulation of Jagged (JAG1) in metastatic PCa compared to localized disease or benign prostate tissue, and a significant association of high JAG1 expression with tumor recurrence. Activation of Notch signaling is associated with EMT in PCa[18], and both normal mouse prostate tissue and human prostate tumor explants display the features of EMT following androgen deprivation[19]. Together, these observations pinpoint a possible role of interplay among Notch-Jagged signaling, AR and EMT in enabling prostate cancer aggressiveness and relapse.

Here, we develop a mechanism-based mathematical model to investigate the emergent dynamics of Notch-EMT-AR signaling axis. Our model predicts that AR can stabilize a hybrid E/M phenotype and thus prevent cells from undergoing a complete EMT, especially at higher levels of Jagged production rates and consequently a dominating Notch-Jagged signaling compared to Notch-Delta signaling. We also observed that the presence of AR can increase the fraction of hybrid E/M cells co-localizing, thus increasing the likelihood of CTC cluster formation. Finally, we observed an association of worse survival of combined expression of AR and JAG1 in clinical samples, demonstrating the aggressive traits associated with hybrid E/M phenotypes.

## 2 Methodology

### 2.1 Mathematical modeling of the GRN circuit

The mathematical formalism of the Notch-EMT-AR axis has been described as the dynamics of the chemical species of the Notch pathway (Notch receptor, NICD, Jagged, Delta), the EMT regulatory module (miR200, miR34, Snail, Zeb) and Androgen Receptor. The interaction between all the species have been illustrated in Figure 1A. Ordinary differential equations have been used to model the temporal dynamics of the species in the circuit. The equations of all the chemical species is presented in Supplementary file. Each of the chemical species has a specific rate of production and decay. Moreover, the processes like transcriptional and translational modulations can affect the production of any species. Information on how these modulations has been implemented in this framework can be found in the Supplementary file. All the parameter values have been tabulated in the supplementary file. Finally, methodological details on the methods implemented to execute the simulations are given in the supplementary file.

**Figure 1.**
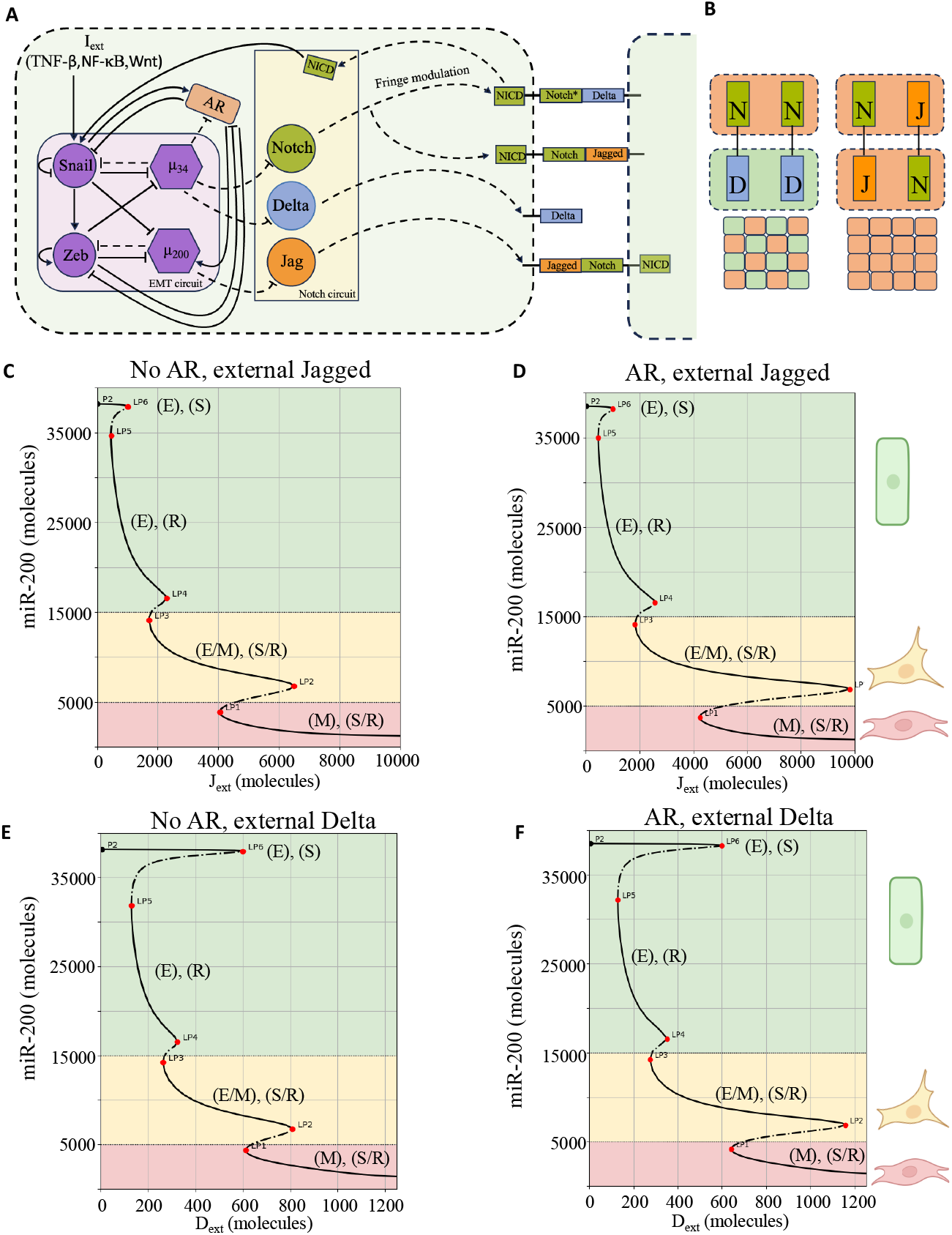
Schematic of the coupled Notch-Delta/Jagged signaling circuit with the EMT circuit along with the bifurcation diagram of miR200 in the presence and absence of Androgen Receptor (AR). (A) Schematic representing the connection between the Notch-Delta/Jagged signaling circuit and the core EMT circuit. Notch and Delta are inhibited by miR34, while Jagged is inhibited by miR200. Snail is activated by NICD. AR inhibits Snail while Snail activates AR. AR-Zeb exhibits a mutual inhibitory loop. AR activates miR200, while miR34 inhibits AR. (B) Cell-cell communication mediated by Notch[N]-Delta[D] gives rise to divergence in cell fate, one cell becoming the sender (S), having a low level of Notch and high Delta, and the other becoming the receiver [R], having a low level of Delta and a high level of Notch. In contrast, when cell-cell communication is mediated through Notch-Jagged[J] interaction, both cells attain a similar sender/receiver phenotype [S/R]. In the context of a multi-cell level, Notch-Delta signaling can exhibit a ‘salt-and-pepper’ pattern. In contrast, Notch-Jagged signaling forms a cluster of cells having a similar sender/receiver phenotype. (C) Bifurcation diagram with respect to level of miR200 as a function of external Jagged concentration (*J*_*ext*_). Here, the level of external Delta is set to 0. (D) Same as C, in presence of AR. (E)Bifurcation diagram with respect to level of miR200 as a function of external Delta concentration (*D*_*ext*_). Here, the level of external Jagged is set to 0. (F)Same as E, in presence of AR. Solid and dotted line represents stable and unstable steady states. The thresholds for the different phenotypes are colored as Epithelial - green, Hybrid Epithelial/Mesenchymal - yellow and Mesenchymal – red with corresponding cartoons. Bifurcations of all the other species are given for the above scenarios as supplementary Figures 1-4.

### 2.2 Numerical solving and analysis

The single-cell and multi-cell formalisms are executed and solved numerically on Python using the numerical library PyDSTool [20]. All plots were generated using GraphPad. Statistical analyses were performed in GraphPad Prism as well, using the unpaired two-tailed Student’s t-test. The significance figures (*P* value) is denoted as: ns (*P* > 0.05), * (*P* ≤ 0.05), ** (*P* ≤ 0.01), *** (*P* ≤ 0.001), **** (*P* ≤ 0.0001).

### 2.3 Transcriptomic Data Analysis

The raw count matrices for bulk RNAseq and scRNAseq dataset was downloaded from NCBI GEO. The bulk RNAseq count matrix was converted to the TPM format. The epithelial, mesenchymal, and partial EMT scores were calculated using ssGSEA function available in GSEApy package. The expression of (genes) and (signatures) were visualized using barplots in the resistant and sensitive group. Statistical t test was performed to assess the significance and performed using the Rstatix package.

The scRNAseq count matrices of three samples were downloaded from GSE168668 and were normalized using the LogNormalize function. They were further integrated together using the RunHarmony function. The Dimensionality reduction was performed on the data using the RunUMAP function. The above preprocessing and their visualization were performed using the Seurat package. The epithelial, mesenchymal, and partial EMT scores were calculated using AUCell package for the (signatures). All bioinformatic analysis was performed on R version 4.3.2.

### 2.4 Survival Analysis

Proggene, a web application for gene expression-based survival analysis, was used to perform survival analysis [21]. The cancer selected was ‘prostate cancer’, and the survival type was ‘death’ and ‘relapse’. The population was divided by the median. Survival analysis was performed for AR, JAG1, and DLL4, where individual and combination graphs were generated.

## 3 Results

### 3.1 AR stabilizes hybrid E/M state at a single cell level

To examine the role of AR on EMT dynamics, we extended the earlier Notch-EMT circuit proposed by Boareto *et al*. [22]to incorporate the regulatory effects of AR on the circuit components [Figure 1A]. First, AR and ZEB mutually inhibit each other [19]. Second, AR suppresses Snail expression [23, 24], while Snail, in turn, activates AR [25, 24]. Third, miR-34 inhibits AR [26, 27], and AR, in turn, activates miR-200 [28, 29]. For simplicity, in this study, we focus exclusively on the wild-type form of AR and exclude the effects of any mutated or alternatively spliced AR isoforms.

Notch signaling is initiated by binding of the trans-membrane Notch receptor of a cell to either Delta or Jagged ligands presented by a neighboring cell. This binding leads to the release of Notch Intra Cellular Domain (NICD) that triggers transcriptional response downstream. At a multicellular level, the nature of the ligand involved determines the resulting phenotypic pattern. Specifically, Notch-Jagged signaling promotes convergent cell fates, wherein both interacting cells adopt a sender/receiver phenotype (medium ligand, medium receptor). In contrast, Notch-Delta signaling results in divergent cell fates, with one cell becoming the sender (high ligand, low receptor) and the other, the receiver (low ligand, high receptor) [22, 30] [Figure 1B]. This property of Notch-Jagged signaling facilitates the formation of clusters of cells with similar phenotypes, and consequently, it can promote the emergence of hybrid E/M cell clusters [22]. This occurs through coupling between the Notch and EMT circuits, where Notch activation induces the EMT-promoting transcription factor Snail. Conversely, the epithelial-state-maintaining microRNAs—miR-34 and miR-200—suppress Notch signaling by downregulating the expression of Notch, Delta, and Jagged through transcriptional repression[31, 22].

First, we studied the dynamics of the Notch-EMT-AR circuit at the intracellular level as a function of the average concentration of the Jagged and Delta ligands which are available on the surface of the neighbor cell and which are implemented as constant levels of external ligands *J*_*ext*_ and *D*_*ext*_. It is well established that the activation of Notch by either Jagged or Delta can induce a complete or partial EMT in context of epithelial cells [22, 32]. Consistently, we observed with the increase in the ligand concentration, regardless of the presence of AR, epithelial cells acquiring a partial or completely mesenchymal state[Figure 1C-F]. Bifurcation plots of all the other species are given as supplementary figures 1-4, in the presence and absence of AR respectively.

At low concentrations of both the ligands in the absence of AR, the cell maintains an epithelial state and can act as either a sender or receiver. With an increase in the concentration of the external ligands, the cell acquires a hybrid E/M phenotype and can act as both sender and receiver. On further increasing the ligand concentrations, the cell acquires a complete mesenchymal phenotype and can act as both sender and receiver. Now, in the presence of AR the overall dynamics are similar, but the range of value of the external ligand for which a hybrid E/M state can exist gets extended (compare the stable fixed points in the yellow region in Figure 1). Hence, a complete mesenchymal progression is hindered. Thus, we conclude that in the presence of AR, the cell requires a much stronger stimulus to reach a complete mesenchymal state at a single cell level.

To test the robustness of the model parameters, we performed sensitivity analysis by varying them by 10%. We found the results to be robust to these variations,although a high sensitivity was observed with the parameters belong to the original EMT circuit [Supplementary Figure 5,6] by Lu *et al*.[33].

In conclusion, our model predicts that at a single-cell level AR can act as a phenotypic stability factor for the hybrid E/M state.

### 3.2 AR alters the tissue-level EMT heterogeneity composition

Following the single-cell level analysis of the effect of AR, we extended our study to tissue level by formulating a two-dimensional lattice of 60 *×* 60 cancer cells capable of intercellular communication via Notch signaling. Through this framework we investigated the behavior of the Notch-EMT and Notch-EMT-AR circuits in a multicellular context. We specifically examined both the relative fractions of epithelial, hybrid E/M, and mesenchymal phenotypes, as well as their spatial distribution within the lattice, under varying production rates of Delta (*g*_*D*_) and Jagged (*g*_*J*_), starting from a randomized initial distribution [Supplementary Figure 7].

We first analyzed the tissue-level dynamics of Notch-EMT and Notch-EMT-AR networks under conditions where cells in the lattice communicated primarily via Jagged-mediated Notch signaling [Figure 2]. All the results were recorded after a transient period of 120 hours (5 days), which corresponds to a typical timescale over which EMT occurs. Beyond this period, phenotypic patterning may be influenced by biophysical processes such as changes in cell morphology and cell migration, which we are not explicitly taking into account in this current study. As previously discussed, Jagged-mediated Notch signaling can promote the formation of clusters of hybrid E/M cells [34]. To explore this aspect, we initially set the production rate of Jagged to a low level (*g*_*J*_ = 45 molecules/h), which induces mild Notch activation and thus a weak EMT response. Under these conditions, AR stabilizes the epithelial phenotype through activation of miR-200 (a promoter of epithelial identity) and inhibiting mesenchymal-promoting transcription factors such as Snail and Zeb [see Figure 1A]. This regulatory influence of AR leads to a reduction in both hybrid E/M and mesenchymal cell populations [compare Figures 2A and 2B]. To quantify these observations, we measured the fraction of each phenotypic state across multiple simulation replicates, accounting for variability introduced by the randomized initial conditions of the lattice. From this analysis, we concluded that at low Jagged production rates (*g*_*J*_ = 45 molecules/h), the presence of AR significantly reduces the proportion of hybrid E/M and mesenchymal cells, thereby increasing the fraction of epithelial cells [Figure 2E].

**Figure 2.**
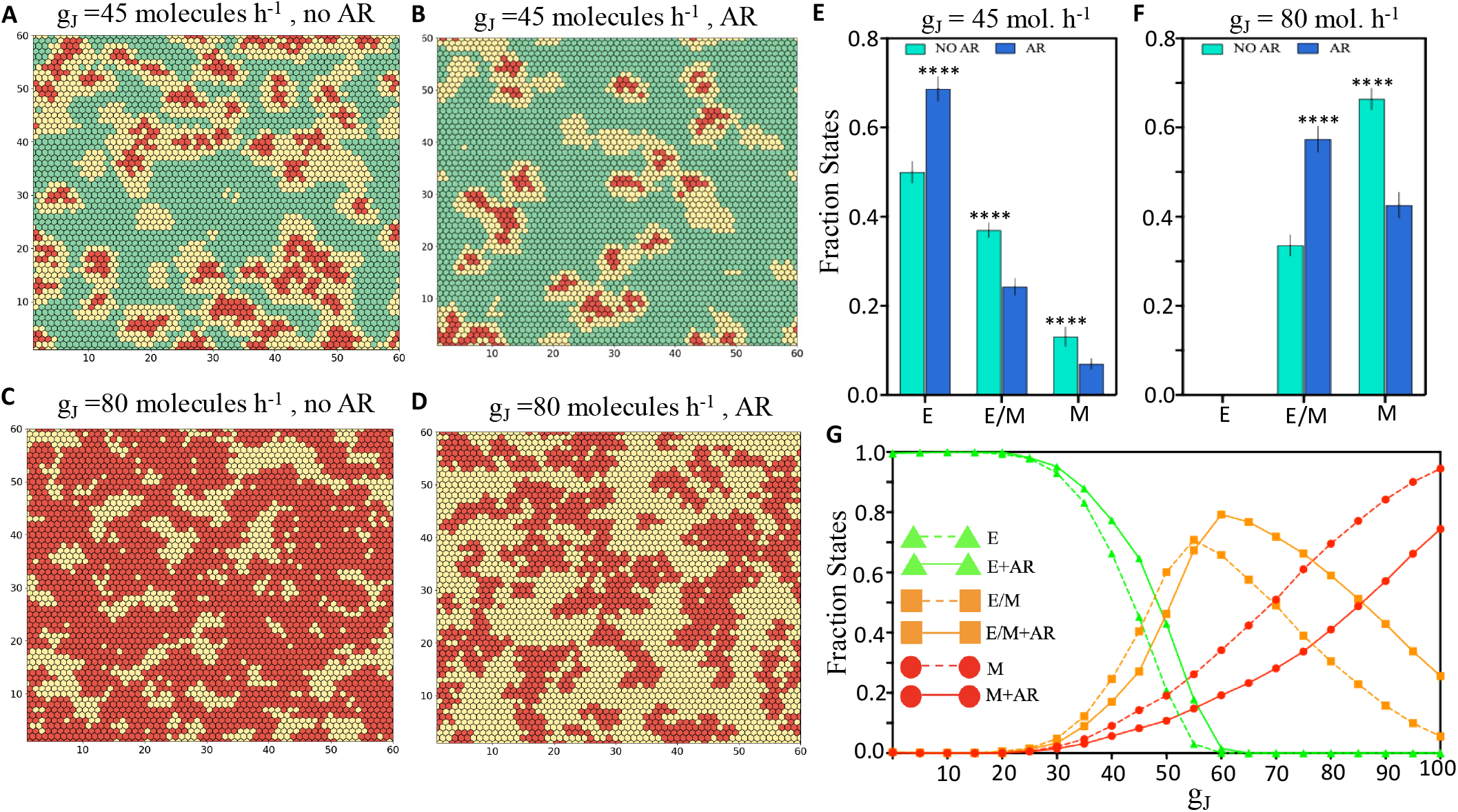
Effect of AR on the tissue patterning of a Jagged dominated Notch signaling lattice. (A) Snapshot of a two dimensional lattice without AR at *g*_*J*_ = 45 molecules/h, (B)Same as (A) in presence of AR, (C) Snapshot of a two dimensional lattice without AR at *g*_*J*_ = 80 molecules/h, (D) Same as (C) in presence of AR, (E) Average fraction of states for *g*_*J*_ = 45 molecules/h, AR decreases both the fraction of cells in hybrid E/M and mesenchymal states,(F) Average fraction of states for *g*_*J*_ = 80 molecules/h, all cells have attained partial EMT, AR reduces number of cells attaining a mesenchymal phenotype, (G) Fraction of E, E/M and M cells as a function of the production rate of Jagged (*g*_*J*_) in a two dimensional lattice in the presence and absence of AR. In the lattice, green represents epithelial state, yellow stands for hybrid state and red stands for mesenchymal state. The production rate of Delta was fixed at *g*_*D*_ = 20 molecules/h for all the simulations, each simulations were started for an initial randomized condition [Supplementary Figure 7]. For each simulation the snapshots and the fraction of states were recorded at 120 hours. For statistical analysis number of replicates were 10. The significance figures are given in the methodology section.

Subsequently, we increased the production rate of Jagged to *g*_*J*_ = 80 molecules/h to investigate tissue-level dynamics under stronger Notch activation. At this higher Jagged production rate, the role of AR appeared to shift. The intracellular levels of NICD increased due to enhanced Notch activation, pushing most cells toward either a hybrid E/M state or a fully mesenchymal state. However, in the presence of AR, its inhibitory effects on the mesenchymal-promoting transcription factors Zeb and Snail prevented cells from undergoing a complete EMT. As a result, a greater fraction of cells remained in a hybrid E/M state [Figures 2C and 2D]. Hence, under this scenario, AR seems to function as a classic phenotypic stability factor (PSF) for the hybrid E/M phenotype, consistent with our observations at the single-cell level analysis. To quantitatively support these observations, we performed multiple simulation replicates and analyzed the distribution of phenotypic states in the presence and absence of AR. Under high Jagged production, AR maintained a significantly higher fraction cells in a hybrid epithelial/mesenchymal state while reducing the population of mesenchymal cells [Figure 2E].

To quantify the spatial co-localization of hybrid E/M cells, we counted, for each hybrid E/M cell, the number of neighboring hybrid E/M cells. Under the hexagonal lattice configuration used in our simulations, each cell has six immediate neighbors. When the production rate of Jagged was set to *g*_*J*_ = 45 molecules/h—corresponding to weak Notch activation—we observed that the presence of AR led to a decrease in the number of hybrid E/M neighbors per hybrid cell. This reduction is a direct consequence of the overall decline in the fraction of hybrid E/M cells [compare Supplementary Figure 8 (left) with 8 (middle)]. However, in the presence of AR, under strong Notch activation (i.e., *g*_*J*_ = 80 molecules/h), the spatial co-localization of hybrid E/M cells was significantly enhanced. This is a direct consequence of both the increase in overall fraction of hybrid E/M cells due to AR and the effect of lateral induction, a phenomenon characteristic of Jagged-mediated Notch signaling that promotes the emergence of neighboring cells with similar phenotypes [34].

Next, we compared the tissue-level patterning dynamics of the Notch-EMT and Notch-EMT-AR networks under conditions where cells primarily communicated via Delta-mediated Notch signaling [Figure 3]. As expected from the lateral inhibition characteristic of Notch-Delta signaling, we observed a ‘salt-and-pepper’ spatial pattern, where neighboring cells tend to acquire opposite phenotypes [34]. Interestingly, for both low and high Delta production rates, there was no significant difference in the overall tissue-level dynamics between the Notch-EMT and Notch-EMT-AR networks, suggesting that AR has limited impact on spatial organization in the context of Delta-dominated signaling.

**Figure 3.**
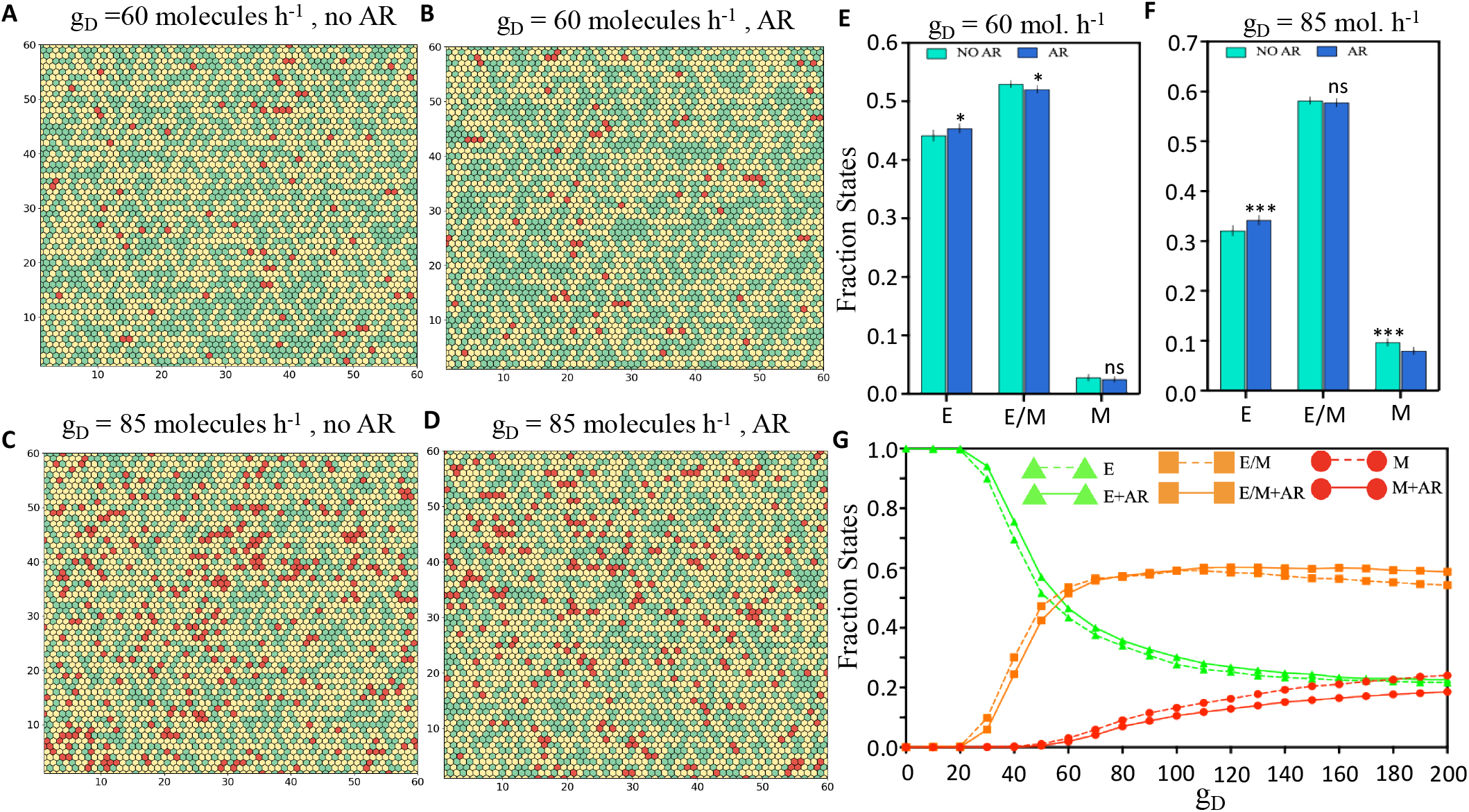
Effect of AR on the tissue patterning of a Delta dominated Notch signaling lattice. (A) Snapshot of a two dimensional lattice without AR at *g*_*D*_ = 60 molecules/h, (B) Same as (A) in presence of AR, (C) Snapshot of a two dimensional lattice without AR at *g*_*D*_ = 85 molecules/h, Same as (C) with AR, (E) Average fraction of states for *g*_*D*_ = 65 molecules/h, presence of AR makes no significant alteration,(F) Average fraction of states for *g*_*D*_ = 85 molecules/h, AR tend to increase epithelial fraction mildly while reducing the mesenchymal population with no significant effect on the hybrid population, (G) Fraction of E, E/M and M cells as a function of the production rate of Delta (*g*_*D*_) in a two dimensional lattice in the presence and absence of AR. In the lattice, reen represents epithelial state, yellow stands for hybrid state and red stands for mesenchymal state. The production rate of Jagged was fixed at *g*_*J*_ = 20 molecules/h for all the simulations, each simulations were started for an initial randomized condition [Supplementary Figure 7]. For each simulation the snapshots and the fraction of states were recorded at 120 hours. For statistical analysis number of replicates were 10. The significance figures are given in the methodology section.

These observations suggest that the presence of AR can stabilize the hybrid E/M phenotype specifically under Notch signaling that is predominantly mediated by Jagged. In contrast, AR has little to no effect on phenotypic dynamics when Notch signaling is driven by Delta. This highlights the potential importance of the Notch-Jagged signaling axis in promoting the pro-oncogenic functions of AR by sustaining the hybrid E/M state.

In the tumor microenvironment, soluble Notch ligands have been reported to activate Notch signaling in a paracrine manner, in addition to the classical juxtacrine mode of activation [35]. To incorporate this biological context, we extended our analysis to study the tissue-level patterning dynamics of the Notch-EMT and Notch-EMT-AR circuits under the conditions where Notch signaling is mediated by soluble ligands(*sL*_*ext*_). Consistent with our previous findings [Figure 2], when the lattice cells communicated through a Jagged-dominated Notch signaling mechanism [Supplementary Figure 11], the presence of AR significantly altered phenotypic distributions depending on the concentration of the external ligand. At a lower concentration (*sL*_*ext*_ = 1000 molecules), AR promoted the epithelial phenotype by stabilizing epithelial markers and inhibiting mesenchymal transcription factors, resulting in a significant decrease in both mesenchymal and hybrid E/M cell populations [Supplementary Figures 11A, B, E]. However, at a higher external ligand concentration (*sL*_*ext*_ = 4000 molecules), AR prevented cells from fully attaining a mesenchymal state. Consequently, significantly increasing the hybrid E/M population[Supplementary Figures 11C, D, F]. These observations suggest that at high levels of soluble ligands, AR enhances the stability of the hybrid E/M state within a Jagged-dominated Notch signaling environment [Supplementary Figure 11G].

When cells in the lattice communicated via Delta-mediated Notch signaling, we observed that, across all concentrations of external ligand, the presence of AR significantly increased the fraction of cells in an epithelial phenotype consequently decreasing population of cells in both hybrid and mesenchymal phenotypes[Supplementary Figures 12E, F, G]. This trend strengthens our hypothesis that Notch signaling mediated exclusively by Jagged, but not the one mediated by Delta, can promote the pro-oncogenic functions of AR by stabilizing hybrid E/M phenotype.

The presence of external ligands also has a significant impact on the temporal dynamics of phenotypic fractions within the system. It can enhance the lifetime of both hybrid E/M and mesenchymal states in the context of both Jagged- and Delta-mediated juxtacrine Notch signaling. In the case of Jagged-mediated signaling, cells in the hybrid E/M phenotype tend to revert to an epithelial state as the system approaches equilibrium [Supplementary Figure 10A]. A similar trend is observed under Delta-dominated signaling, where hybrid E/M cells also transition back to an epithelial state over time [Supplementary Figure 10B]. However, the presence of external ligands can stabilize the hybrid E/M phenotype, thereby extending its temporal persistence and supporting the maintenance of hy-brid E/M clusters within the lattice under Notch-Jagged signaling [Supplementary Figures 10C,D]. These observations suggest that soluble ligands in the microenvironment can further enhance the stability and persistence of the hybrid E/M phenotype.

It is important to note that signaling mediated by soluble ligands, whether soluble Delta or soluble Jagged, is mechanistically distinct from the intracellular feedback loops between Notch-Jagged and Notch-Delta, which are primarily responsible for spatial patterning. Therefore, whether Notch signaling is activated by soluble Delta or soluble Jagged does not affect the outcome. In this context, the cell function solely as “receiver” or “target” of the soluble ligands. These observations suggest that AR exerts a comparable influence on the dynamics of the system, regardless of whether Notch signaling is driven by soluble Delta or soluble Jagged.

Together, these simulation results over a multi-cell lattice setup highlights how AR signaling can alter the spatial patterning and heterogeneity in terms of EMT composition at a tissue-level. Overall, it increases the fraction of hybrid E/M cells in a population only under conditions of dominant Notch-Jagged signaling.

### 3.3 AR attenuates epithelial-mesenchymal transition induced by external EMT signal

Next, we studied the effect of AR in the presence of external EMT inducing signal like TGF-*β*, NF-*κ*B, Wnt, that directly activates Snail [Figure 1A]. We first analyzed the dynamics of Notch-EMT and Notch-EMT-AR circuits in a lattice of cells communicating via Jagged dominated Notch signaling [Figure 4A,B]. Higher strength of the EMT signal(*I*_*ext*_) can push a significant amount of cells into a completely mesenchymal state [Figure 4G]. While in the presence of AR for a fixed rate of the external signal, the amount of cells in a complete mesenchymal state is significantly reduced [Figure 4G,E]. This observation is consistent with our previous single-cell simulation results that AR acts as a brake in the progress towards a complete mesenchymal state and hence a higher rate of external EMT inducing signal is required to push more cells towards a mesenchymal state. One striking feature to notice here is that the fraction of hybrid E/M cells were minimum under this scenario [Figure 4E, G]. This can potentially explain the experimental observation that knockdown of AR can promote EMT in prostate cells [36, 23].

**Figure 4.**
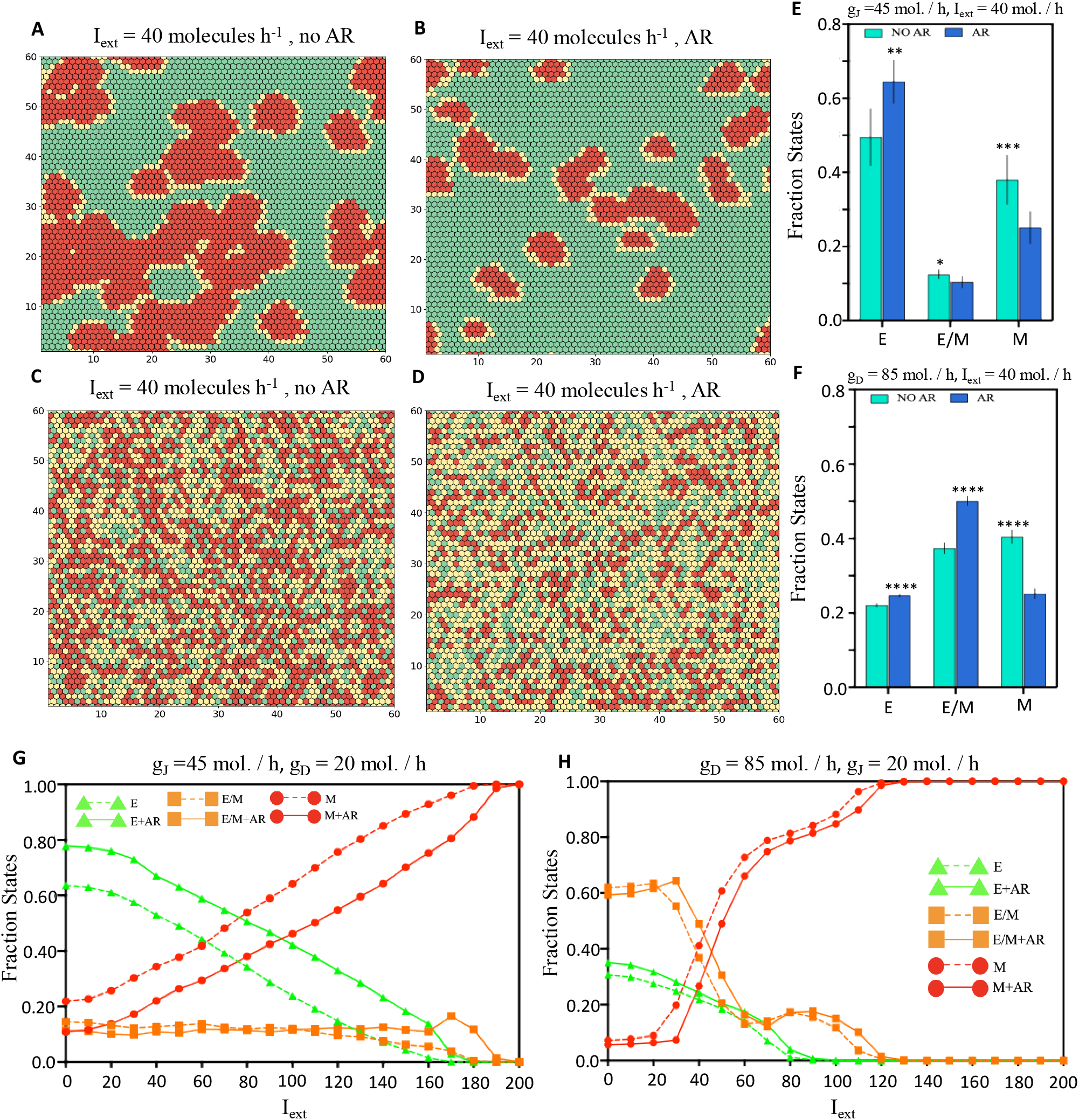
Effect of AR on the tissue patterning under the influence of external EMT signal in case of Jagged dominated and Delta dominated Notch signaling lattices. (A) Snapshot of a two dimensional lattice communicating through Jagged dominated Notch without AR under external EMT signal of *I*_*ext*_ = 40 molecules/h. (B) Same as (A) in presence of AR. (c) Snapshot of a two dimensional lattice communicating through Delta dominated Notch without AR under external EMT signal of *I*_*ext*_ = 40 molecules/h. (D) Same as (C) in presence of AR. (E) Fraction of states plot for the case depicted in (A) and (B). (F) Same as (E) for (C) and (D). (G) Plot of the fraction of states with respect to the external EMT signal rate under Jagged dominated Notch signaling. (H) Same as (G) for a Delta dominated signaling lattice. In both the Jagged dominated and Delta dominated scenario the production rate of the other ligand was fixed at 20 molecules/h. The simulation in (A) and (B) was started from Figure 2A, B as the initial configuration for the case with and without AR respectively. Similarly (C) and (D) started from Figure 3C,D as the initial configuration. The total number of replicates taken for each scenario was 10. The significance figures

Next, we examined the scenario in which cells within the lattice primarily communicated through Delta-mediated Notch signaling. Consistent with observations in the Jagged-dominated signaling case, the presence of AR led to a reduction in the fraction of cells exhibiting a mesenchymal phenotype, although this effect was less pronounced compared to the Jagged-dominated lattice. Notably, unlike the Jagged-dominated case, AR seems to also enhance the proportion of cells having a hybrid epithelial/mesenchymal phenotype over some specific range of external signal (*I*_*ext*_).

Finally, to assess the temporal stability conferred by AR to the hybrid E/M phenotype, we initialized a lattice composed exclusively of hybrid E/M cells by randomly sampling parameter values from within the ranges that permit existence of the hybrid E/M state [Supplementary Figure 13]. EMT was then induced across the lattice using an external EMT-inducing signal which activates Snail, driving the cells toward a fully mesenchymal phenotype. We defined the half-life of the hybrid state as the time point at which the ratio of hybrid E/M to mesenchymal cells reached 1:1 and recorded this metric. In the absence of AR, the average half-life of hybrid cells was approximately 115 hours. However, in the presence of AR, this half-life was extended to 157 hours [Supplementary Figure 13C]. These results suggest that AR can significantly enhance the temporal stability of the hybrid E/M state, thereby increasing the probability of forming circulating tumor cell (CTC) clusters through the spatial localization of these hybrid E/M cells.

### 3.4 AR and JAG1 are co-expressed in androgen-resistant prostate cancer cohorts in in vitro and patient samples

We next investigated whether the predictions from our mechanism-based mathematical model can be supported using publicly available experimental data. Thus, we obtained bulk and single-cell transcriptomic data consisting of both androgen-sensitive and resistant samples from in vitro experiments and patient tumors, respectively, to assess how commonly are AR and JAG1 signlaing co-occuring and what is the implication of this co-occurence in androgen resistance.

The bulk RNAseq data is taken from dataset GSE123379 and consists of enzalutamide-sensitive and resistant C4-2 prostate cancer adenocarcinoma cells [16]. This study demonstrated that Notch signalling contributed to enzalutamide resistance in C4-2 cell line, thereby making it a suitable to analyse the AR-JAG1 interplay. The normalized gene expression of AR, JAG1 and DLL4 were visualized using a barplot, and we observed that AR and JAG1 were upregulated in the enzalutamide resistant cohort, whereas DLL4 was downregulated [Fig 5A]. This trend suggests that resistance to enzalutamide in prostate cancer could be mediated through AR and JAG1-mediated Notch signaling axis, as both NOTCH2 and NOTCH3 expression is also higher in resistant cells [Supplementary Figure 16]. A similar analysis was conducted using single-cell RNA-seq of castration-sensitive and resistant prostate cancer (CSPC and CRPC, respectively) patient tumor samples obtained from dataset GSE206962. Here, castration resistance was acquired by Androgen Deprivation Therapy(ADT) and treatment with androgen receptor antagonists like enzalutamide. Comparable findings were noted in the case of single-cell data as well, especially in the resistant1 cohort [Fig 5B-E]. These observations highlight that resistance to AR targeting agents such as enzalutamide often displays a co-occurence of upregulation of AR and Notch-Jagged signaling that may stabilize a hybrid E/M phenotype, therefore aggravating tumor progression, as predicted by our mechanistic model earlier.

**Figure 5.**
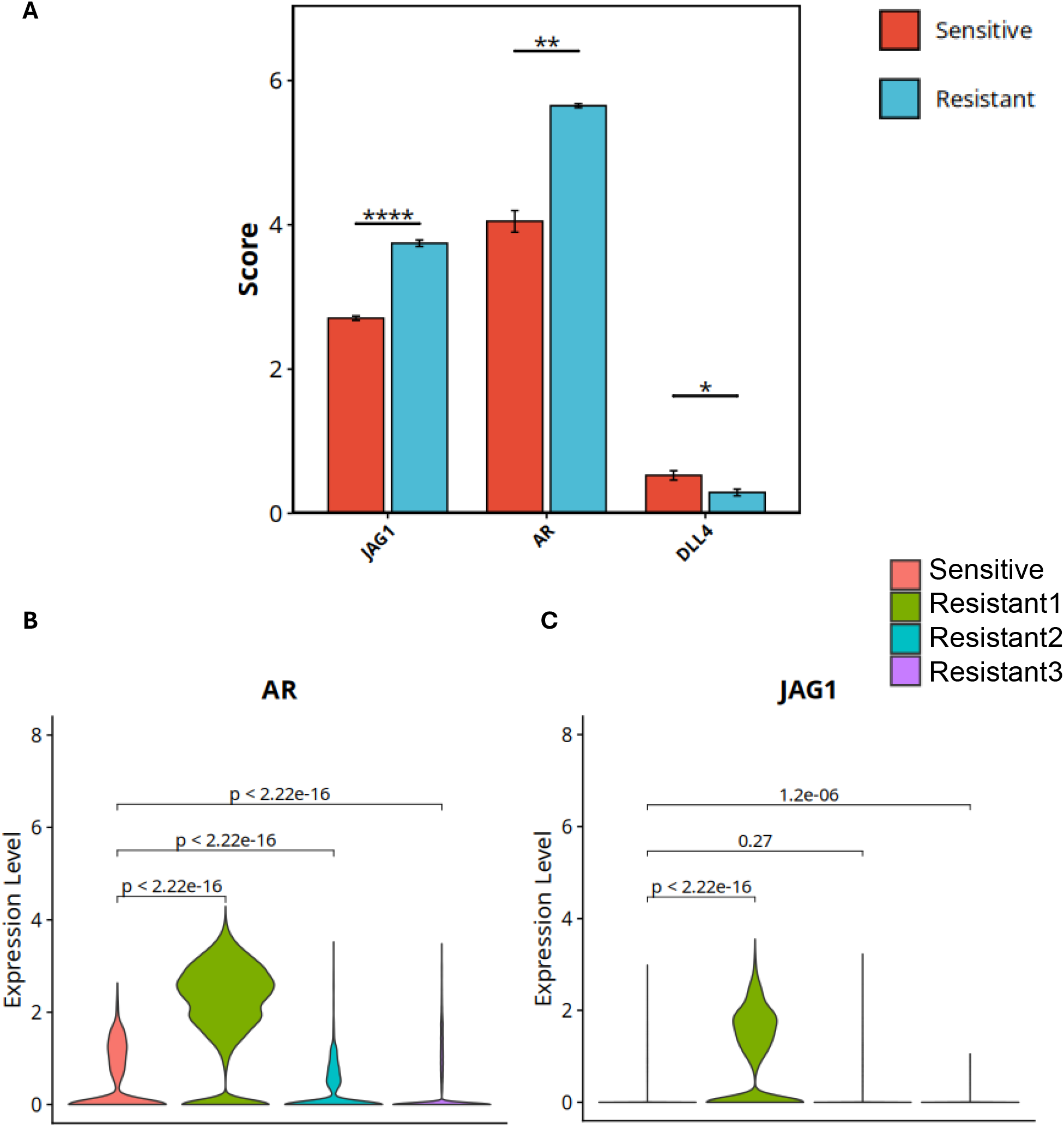
Co-expression of AR and JAG1 in in-vitro and patient data. (A) Boxplots showing expression of AR, JAG1 and DLL4 gene expression for sensitive and resistant cell lines from GSE123379 [16].Unpaired t-test was performed and the p-value is reported(*: p¡0.05,**: p¡0.01 and ****:p¡0.000. (B)-(C) Violin plots depicting the gene expression of AR and JAG1 for four patient single cell RNAseq data (1 Castration sensitive and 3 Castration resistant PCa from GSE206962). The Wilcoxon test was performed and the p values are reported in the figure.

### 3.5 AR and JAG1 co-expression is correlated with worse prognosis in prostate cancer patients

Finally, we look at clinical data to evaluate the effect of AR-JAG1-mediated signalling on patient survival and cancer progression. For this assessment, we used PROGgene to generate Kaplan-Meier curves for high gene expression vs low gene expression of the three genes-AR, JAG1 and DLL4, both individually and combined. Our model predictions that AR can stabilize hybrid E/M state under the influence of Notch-Jagged signaling, but not under Notch-Delta signaling. Thus, we investigated the impact of measuring survival when both AR and JAG1 are upregulated vs. when both AR and DLL4 are upregulated. We observed that only AR expression does not have a significant impact on the overall survival. Although, JAG1 high expression corresponds to worse survival, the combined expression of JAG1 and AR has even worse survival probability, as indicated by a higher hazard ratio[Fig 6A]. Similarly, high AR expression does not correspond to worse relapse free survival (RFS), but high JAG1 has a significant effect on RFS. Additionally,the combined expression of JAG1 and AR has worse probability of cancer progression, as indicated by a higher hazard ratio [Fig 6B,C]. It is also worth noting that high expression of DLL4 alone, and in combination with high AR is not correlated, with worse prognosis [Supplementary Figure 17].

**Figure 6.**
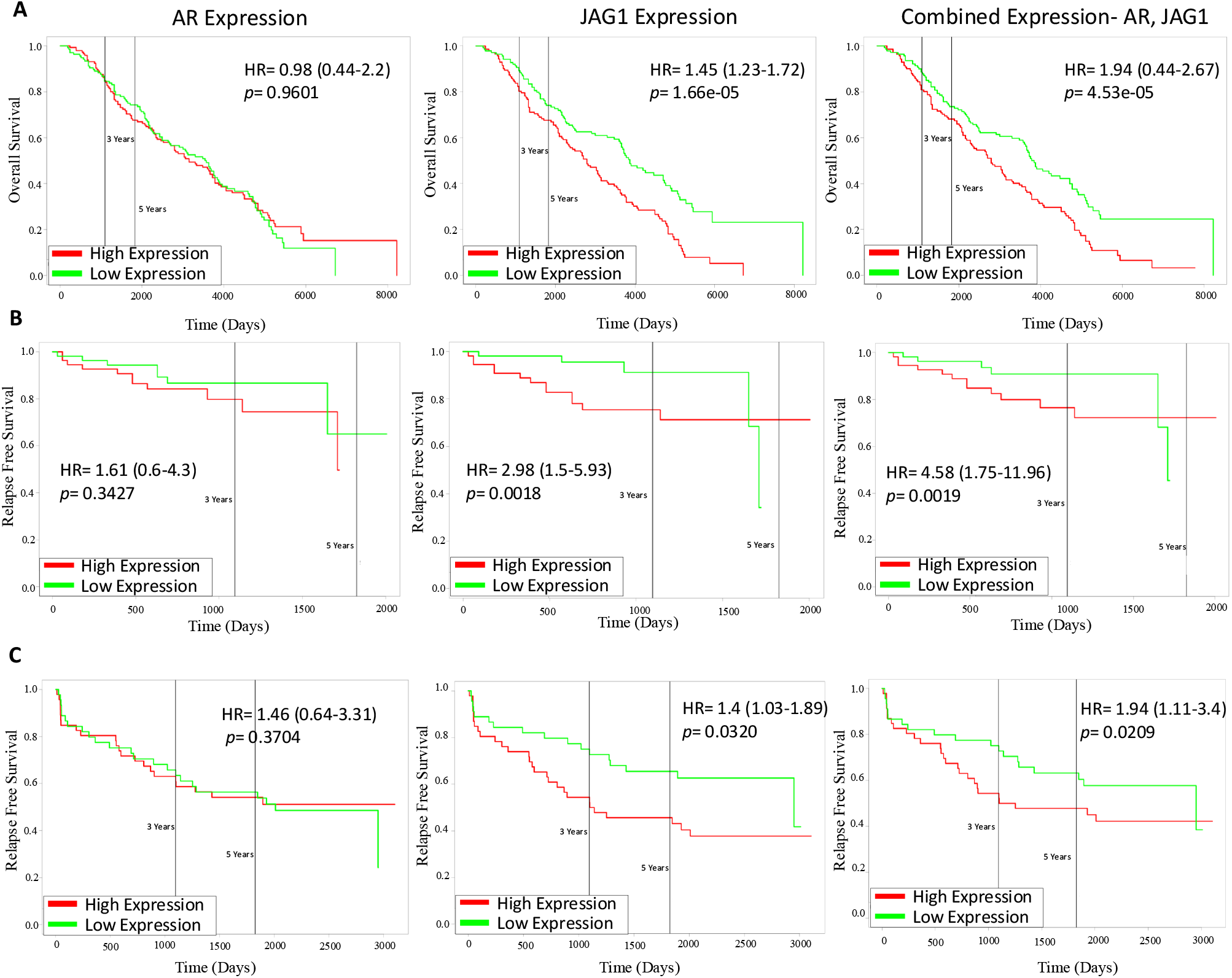
Survival analysis for co-expression of AR and JAG1 in several PCa patient cohorts. (A) Kaplan–Meier curves for overall survival (OS) for AR high vs low expression (left), JAG1 high vs low expression (centre), and Both AR and JAG1 combined high vs low expression (right) in prostate cancer patient data from GSE16560 . Reported p-values are based on a log-rank test a indicating significant difference in survival. Same as (A) but for (B), Relapse Free Survival in prostate cancer patient data from GSE70768, and (C) Relapse Free Survival in prostate cancer patient data from GSE70769.

## 4 Discussion

Recent in vitro and in vivo evidence have demonstrated the highly tumorigenic, metastatic, and therapy-evasive traits of the hybrid E/M phenotypes [37, 5, 38]. Thus, identifying the molecular signatures and therapeutic vulnerabilities of these phenotypes is crucial to develop anti-metastatic therapies. Previous work has suggested many possible ‘phenotypic stability factors’ for hybrid E/M phenotypes, such as miR-129 [39], Connexin 43 [40], NRF2 [41], P4HA2[42], ITGB4 [43] and calcium signaling including the role of NFATc [44, 45]. Our work suggests that the interplay among intracellular EMT and AR pathways, and Notch-Jagged cell-cell communication can also stabilize the hybrid E/M phenotypes, thus improving our understanding of how non-cell autonomous factors can control tumor progression.

Extending our simulation results, the worse patient survival trends observed upon co-expression of AR and JAG1 can be potentially attributed to their joint role in stabilizing the hybrid E/M phenotype(s). Growing clinical evidence endorses the association of hybrid E/M phenotypes with worse outcomes in breast cancer[46], bladder cancer[47], colorectal cancer[48]and head and neck cancer[49]. However, the molecular mediators and clinical implications of hybrid E/M phenotypes in prostate cancer are relatively less studied. Therefore, our results highlight how context-specific mechanistic models of EMT can help identify new potential combinations of biomarkers to assess disease aggressiveness.

Our current framework has several limitations. First, we only consider the role of full-length wild-type AR and not any of its mutated or truncated versions, for instance, AR-v7 that has been associated with resistance to inhibitors of AR signaling such as enzalutamide[13]. Second, other mediators of EMT or MET that are known to interact with AR are missing in our regulatory network, such as AR activating SLUG[50] or feedback loop between AR and GRHL2 [51]. Incorporating these nodes and links in our framework can be helpful to decode how these additional interactions shape the landscape of epithelial-mesenchymal plasticity in prostate cancer and associated traits like therapy resistance. Third, our simulations of biochemical regulatory networks have been conducted in a static cell-based lattice, unlike agent-based or multi-scale setups including mechanical attributes of interactions with the extra-cellular matrix (ECM)[52, 53, 54]. Despite these limitations, our work exemplifies how juxtacrine signaling can shape the patterns of phenotypic plasticity and heterogeneity in a tumor cell population.

To conclude, our mathematical modeling framework predicts that AR can stabilize a hybrid E/M phenotype and thus prevent a complete EMT, particularly under the scenarios of predominant Notch-Jagged signaling relative to Notch-Delta. The association of the co-expression of AR and JAG1 with castration resistance and worse clinical outcomes strengthens our predictions, given the reports on the association of the hybrid E/M phenotype(s) as being the ‘fittest’ for metastasis [55]. Our work also suggests potential therapeutic targets to limit metastasis, drug resistance and consequent relapse.

## 5 Code Availability

All the codes used in the current study are available under the following GitHub repository: https://github.com/guhasouvik/Notch-Androgen_Receptor_Project

## 6 Acknowledgements

MKJ was supported by Param Hansa Philanthropies. SR was supported by Axis Bank Graduate Fellowship, Axis Bank Center for Applied Mathematics and Computing, IISc Bangalore. BT was supported by IISc Institute of Eminence (IoE) postdoctoral felowship.

## 7 Author contributions

MKJ conceptualized and supervised the study, and obtained funding. BT contributed to supervising the study, and to data analysis. SG and SR contributed to performing research and data analysis, and to writing the first draft of the manuscript.

