## Supplementary Figures and Text for "Androgen receptor activation stabilizes a hybrid epithelial/mesenchymal phenotype in presence of Notch-Jagged signaling"

#### 1 EMT-Notch-AR Circuit

The circuit in Figure 1a has been realized by generalization of the mathematical model proposed by Boareto et al. [1], which combines the mathematical model of the Notch-Delta/Jagged cell signaling model developed by Boareto, Jolly et al.[2, 3] with the mathematical framework of the core EMT regulatory circuit proposed by Lu et al.[4]. The system dynamics of the Notch-Delta/Jagged circuit is governed by the following Ordinary Differential Equations.

$$\frac{dN}{dt} = k_P g_N H^S(I,N) P_l(\mu_{34},2) - N[(k_{cD}D + k_{cJ}J) + (k_{tD}D_{ext} + k_{tJ}J_{ext})] - \gamma N \tag{1}$$

$$\frac{dD}{dt} = k_P g_D H^S(I,D) P_l(\mu_{34},3) - D(k_{cD}N + k_{tD}N_{ext}) - \gamma D \tag{2}$$

$$\frac{dJ}{dt} = k_P g_J H^S(I,J) P_l(\mu_{200},5) - J(k_{cJ}N + k_{tJ}N_{ext}) - \gamma J \tag{3}$$

$$\frac{dI}{dt} = N(k_{tD}D_{ext} + k_{tJ}J_{ext}) - \gamma I \tag{4}$$

Where $\gamma$ represents the decay rate of Notch, Delta and Jagged, and $\gamma_I$ represents the degradation rate of NICD. The cis-inhibition rates are defined as $k_{cD}$ and $k_{cJ}$ for Notch-Delta and Notch-Jagged respectively. The trans-activation rate is defined as $k_{tD}$ and $k_{tJ}$ for Notch-Delta and Notch-Jagged respectively. The trans activations are functions of the amount of NICD (I) and is defined as $k_t(I) = k_t H^S(I, \lambda_F)$, where Fringe effect is included as $\lambda_F$, where $\lambda_{F,J}$ and $\lambda_{F,D}$ stands for the fold-change for activation by Jagged and Delta respectively. $k_p$ accounts for the translation rate which is assumed to be equal for all the proteins. The transcription rates are $g_N$, $g_D$ and $g_J$ for Notch, Delta and Jagged respectively. The level of protein available for binding either as membrane bound protein of neighbouring cell or present in soluble form is given by $N_{ext}$, $D_{ext}$ and $J_{ext}$ respectively. It was assumed that the modulation of the production rates are implemented the shifted Hill function defined as follows:

$$H^{(S)}(A,B) = \lambda_{A,B}\frac{(A/A_{0,B})^n}{1+(A/A_{0,B})^n} + \frac{1}{1+(A/A_{0,B})^n} \tag{5}$$

where $A_{0,B}$ implies the half-maximal concentration of A with respect to B and $\lambda_{A,B}$ signifies the fold change of the production of B due to A. $\lambda_{A,B} > 1$ implies an activation of B by A and $\lambda_{A,B} < 1$ implies an inhibition of B by A.

The Notch receptor and ligands are also degraded through the post translational regulation due to the binding of the micro-RNAs (miR200, miR34) which has been implemented through the function $P_l(\mu, n)$, which represents the post translational inhibition by a microRNA of type $\mu$ with

$n$ number of corresponding available binding sites. The subsequent degradation of the microRNA complex after binding to the protein mRNA has been implemented through the function $P_y(\mu, n)$. The functions defining the microRNA post translational interactions has been adopted from the work of Lu et al. [4] on microRNA- Based-Chimeric circuits and are defined as follows:

$$P_l(\mu, n) = \frac{L(\mu, n)}{Y_m(\mu, n) + k_m} \tag{6}$$

$$P_y(\mu, n) = \frac{Y_\mu(\mu, n)}{Y(\mu, n) + k_m} \tag{7}$$

where $L(\mu, n)$ represents the total translation rate of the protein, $Y_m(\mu, n)$ implies the net active degradation rate of the mRNA of the protein and $Y_\mu(\mu, n)$ is the net degradation rate of the microRNA. The functions are defined as follows:

$$L(\mu, n) = \sum_{i=0}^{n} l_i C_i^n M_i^n(\mu) \tag{8}$$

$$Y_m(\mu, n) = \sum_{i=0}^{n} \gamma_{mi} C_i^n M_i^n(\mu) \tag{9}$$

$$Y_m(\mu, n) = \sum_{i=0}^{n} \gamma_{\mu i} C_i^n M_i^n(\mu) \tag{10}$$

where $l_i$, $\gamma_{mi}$, $\gamma_{\mu i}$ are the rates for the scenario when $i$ microRNAs are bound to the protein. $C_i^n$ stands for the number of possible arrangements of i molecules of microRNA in n-binding sites:

$$C_i^n = \frac{n!}{i!(n-i)!} \tag{11}$$

$$M_i^n = \frac{(\frac{\mu}{\mu_0})^i}{(1 + \frac{\mu}{\mu_0})^n} \tag{12}$$

where $\mu_0$ represents the threshold fo the concentration of microRNA. Kindly refer to the supplementary sheet of Lu et al. [4] for the detailed version of the above formalism.

The dynamics of the EMT-AR circuit is defined by the following Ordinary Differential Equations.

$$\frac{dS}{dt} = k_{pgs} g_s H^S(S, S) H^S(AR, S) H^S(I, S) H^S(I_{ext}, S) P_I(\mu_{34}, 2) - \gamma_s S \tag{13}$$

$$\frac{dZ}{dt} = k_p g_Z H^S(AR, Z) H^S(Z, Z) H^S(S, Z) P_I(\mu_{200}, 6) - \gamma_Z Z \quad \text{(S8)} \tag{14}$$

$$\begin{aligned}\frac{d\mu_{200}}{dt} &= k_p g_{\mu_{200}} H^S(AR, \mu_{200}) H^S(Z, \mu_{200}) H^S(S, \mu_{200}) - g_Z H^S(Z, Z) H^S(S, Z) P_y(\mu_{200}, 6) \\ &\quad - g_J H^S(I, J) P_y(\mu_{200}, 5) - \gamma_{\mu_{200}} \mu_{200}\end{aligned} \tag{15}$$

$$\frac{d\mu_{34}}{dt} = k_p g_{\mu_{34}} H^S(S, \mu_{34}) H^S(Z, \mu_{34}) - g_S H^S(S, S) H^S(I, S) H^S(I_{\text{ext}}, S) P_y(\mu_{34}, 2)$$
$$- g_D H^S(I, D) P_y(\mu_{34}, 3) - g_N H^S(I, N) P_y(\mu_{34}, 2) - \gamma_{\mu_{34}} \mu_{34} \quad (16)$$

where $g_{\mu_{200}}$, $g_{\mu_{34}}$, $g_z$ and $g_s$ represents the transcription rate of miR200, miR34, Zeb and Snail, respectively; $\gamma_{\mu_{200}}$, $\gamma_{\mu_{34}}$, $\gamma_z$ and $\gamma_s$ represents the degradation rate of miR200, miR34, Zeb and Snail, respectively. The functions $H^{(S)}(A, B)$, $P_l(\mu, n)$, $P_y(\mu, n)$ has already been discussed previously.

#### 1.1 miR-34 inhibition on AR

The inhibition by miR-34 on AR has been implemented using the framework of mRNA based chimeric circuits as discussed previously taking into account the mRNA ($m_{AR}$) and protein ($AR$) level dynamics of AR. The mRNA dynamics equation is written as:

$$\frac{dm_{AR}}{dt} = g_{AR} H^S(Z, AR) H^S(S, AR) - m_{AR} Y_m(\mu_{34}, 2) - \gamma_{m_{AR}} m_{AR} \quad (17)$$

The protein-level dynamics is given as:

$$\frac{dAR}{dt} = k_P m_{AR} L(\mu_{34}, 2) - \gamma_{AR} \quad (18)$$

Here $\gamma_{m_{AR}}$ and $\gamma_{AR}$ represents the degradation rate of AR mRNA and protein respectively and $k_P$ is the translation rate. The functions $Y_m(\mu_{34}, 2)$ and $L(\mu_{34}, 2)$ implemens the mRNA degradation and translation rate for the binding of $\mu_{34}$ to the mRNA $m_{AR}$, where n=2 is the number of binding sites for $\mu_{34}$ on $m_{AR}$. We assume the translation dynamics to be much faster than all the other processes, hence under quarsi stable approximation:

$$\frac{dm_{AR}}{dt} = 0, \quad (19)$$

Then,

$$m_{AR} = \frac{g_{AR} H^S(Z, AR) H^S(S, AR)}{Y_m(\mu_{34}, 2) + \gamma_{m_{AR}}} \quad (20)$$

Hence, the protein level equation 18 becomes:

$$\frac{dAR}{dt} = k_P g_{AR} H^S(Z, AR) H^S(S, AR) \frac{L(\mu_{34}, 2)}{Y_m(\mu_{34}, 2) + \gamma_{m_{AR}}} - \gamma_{AR} AR \quad (21)$$

Using the definition 6

$$\frac{dAR}{dt} = k_P g_{AR} H^S(Z, AR) H^S(S, AR) P_l(\mu_{34}, 2) - \gamma_{AR} AR \quad (22)$$

The updated equation for the dynamics of miR-34 16 which implements the degradation of miR-34 due to the active binding which is implemented by the term $g_{AR} H^S(Z, AR) H^S(S, AR) P_y(\mu_{34}, 2)$ becomes:

$$\frac{d\mu_{34}}{dt} = k_p g_{\mu_{34}} H^S(S, \mu_{34}) H^S(Z, \mu_{34}) - g_S H^S(S, S) H^S(I, S) H^S(I_{\text{ext}}, S) P_y(\mu_{34}, 2)$$
$$- g_D H^S(I, D) P_y(\mu_{34}, 3) - g_N H^S(I, N) P_y(\mu_{34}, 2)$$
$$- g_{AR} H^S(Z, AR) H^S(S, AR) P_y(\mu_{34}, 2) - \gamma_{\mu_{34}} \mu_{34} \quad (23)$$

#### 2 Parameter Estimation

The Parameter values used in the Notch-Delta/Jagged model has been borrowed from the original model developed by Boareto et al. [1] and has been presented in Table 2. Similarly the parameters implemented in the core EMT module has been taken from the original model of Lu et al. [5] and has been presented in Table 3. Please refer to the original articles mentioned above to know in details about the estimation of the parameters. The principle approach followed was as follows:

- The rate of degradation were estimated from literature.
- The Hill fold change were estimated from experiments like Western Blotting involving the the knockdown of the gene or microRNA of interest. The Hill coefficient and the thresholds were estimated from the relative expressions of the inducers or inhibitors or in absence of any literature references were kept at a level to cause tangible effect.
- The cis-inhibition and trans-activation coefficients for the receptor-ligand dynamics of the Notch circuit were set with the assumption that the Notch dynamics is relatively slower than the EMT dynamics.

The parameters associated with AR are presented in Table 4. The degradation rate of AR was kept same as the degradation rates of the other proteins present in the model. The production rate of was estimated to maintain an equilibrium reference level of $10^4$ AR molecules in the system in the absence of any interactions. The Hill fold-changes were estimated from the experimenta papers referred in the main text. The rate of translation was kept same as all the other proteins. The rate of production of the mRNA was calculated in the absence of any interactions from other species.

$$\frac{dm_{AR}}{dt} = g_{AR} - \gamma_{m_{AR}} m_{AR} \tag{24}$$

Assuming quarsi-stable state:

$$m_{AR} = \frac{g_{AR}}{\gamma_{m_{AR}}} \tag{25}$$

Hence, the protein level equation can be written as:

$$\frac{dAR}{dt} = k_p m_{AR} - \gamma_{AR} AR \tag{26}$$

or

$$\frac{dAR}{dt} = k_p \frac{g_{AR}}{\gamma_{m_{AR}}} - \gamma_{AR} AR \Longleftrightarrow \frac{dAR}{dt} = (AR)_0 - \gamma_{AR} AR \tag{27}$$

where $(AR)_0$ is the production rate. Hence to keep $(AR)_0 = 10000$ molecules/h. The production rate of $m_{AR}$ should be 10 $mRNA/h$ as given by the following expression:

$$g_{AR} = \frac{(AR)_0 \gamma_{m_{AR}}}{k_p} \tag{28}$$

where $\gamma_{m_{AR}}$=0.1 molecules/hour (same as the degradation rates of the other ligands and transcription factors) and $k_p$= 100 molcecules/hour.

Table 1: Parameters used in the Notch Circuit. (*) were varied in the multi-cell simulations.

| **Parameter Group** | **Parameters** | **Values** | **Dimensions** |
|---|---|---|---|
| Degradation | $\gamma_N$, $\gamma_D$, $\gamma_J$, $\gamma_I$ | 0.1, 0.1, 0.1, 0.5, 0.1 | $h^{-1}$ |
| Production | $g_N$, $g_D$, $g_J$ | 8, 20(*), 70(*) | mRNA $h^{-1}$ |
| Hill Coefficients | $n_{I,N}$, $n_{I,D}$, $n_{I,J}$ | 2, 2, 5 | Dimensionless |
| Hill Fold-Changes | $\lambda_{I,N}$, $\lambda_{I,D}$, $\lambda_{I,J}$ | 2, 0, 2 | Dimensionless |
| Hill Thresholds | $I_{0,N}$, $I_{0,D}$, $I_{O,J}$ | 200, 200 | Molecules |
| Binding Rates | $k_t$, $k_c$ | $10^-4$, $10^{-5}$ | $h^{-1}$ · Molecules$^{-1}$ |
| Fringe Parameters | $n_F$, $\lambda_{F,D}$, $\lambda_{F,J}$ | 1.0, 3.0(*), 0.3(*) | Dimensionless |

Table 2: Parameters used in the EMT circuit. $K = 1000$ molecules.

| **Parameter Group** | **Parameters** | **Values** | **Dimensions** |
|---|---|---|---|
| Degradation | $\gamma_S$, $\gamma_Z$, $\gamma_{\mu 34}$, $\gamma_{\mu 200}$ | 0.1, 0.1, 0.5, 0.5 | $h^{-1}$ |
| Production | $g_S$, $g_Z$, $g_{\mu 34}$, $g_{\mu 200}$ | 90, 11, 1350, 2100 | mRNA · $h^{-1}$ |
| Hill Coefficient | $n_{Z,\mu 34}$, $n_{Z,\mu 200}$, $n_{S,\mu 34}$, $n_{S,\mu 200}$, $n_{Z,Z}$, $n_{S,S}$, $n_{S,Z}$, $n_{I,S}$, $n_{I_{ext},S}$, $n_{Z,AR}$, $n_{S,AR}$ | 2, 3, 1, 2, 2, 1, 1, 2, 2, 2, 2 | Dimensionless |
| Hill Fold-Change | $\lambda_{Z,\mu 34}$, $\lambda_{Z,\mu 200}$, $\lambda_{S,\mu 34}$, $\lambda_{S,\mu 200}$, $\lambda_{Z,Z}$, $\lambda_{S,S}$, $\lambda_{S,Z}$, $\lambda_{I_{ext},S}$, $\lambda_{S,\text{ext}}$, $\lambda_{Z,AR}$, $\lambda_{S,AR}$ | 0.2, 0.1, 0.1, 0.1, 7.5, 0.1, 10, 6.5, 6.5, 0.5, 2 | Dimensionless |
| Hill Threshold | $Z_{0,\mu 34}$, $Z_{0,\mu 200}$, $S_{0,\mu 34}$, $S_{0,\mu 200}$, $Z_{0,Z}$, $S_{0,S}$, $S_{0,Z}$, $I_{0,S}$, $I_{ext,S}$, $Z_{0,AR}$, $S_{0,AR}$ | 600K, 220K, 300K, 180K, 25K, 200K, 180K, 300, 300, 600K, 450K | Molecules |

Table 3: Parameters of translation, mRNA degradation and micro-RNA degradation upon protein-micro-RNA interaction.

| **Parameter group** | **Parameter** | **Value** | **Dimensions** |
|---|---|---|---|
| Translation rate | $l_i$ | 1.0, 0.6, 0.3, 0.1, 0.05, 0.05, 0.05 | $h^{-1}$ |
| mRNA degradation rate | $\gamma_{mi}$ | 0, 0.04, 0.2, 1.0, 1.0, 1.0, 1.0 | $h^{-1}$ |
| micro-RNA degradation rate | $\gamma_{\mu i}$ | 0, 0.005, 0.05, 0.5, 0.5, 0.5, 0.5 | $h^{-1}$ |

Table 4: Parameters used in the respect to Androgen Receptor.$K = 1000$ molecules.

| **Parameter Group** | **Parameters** | **Values** | **Dimensions** |
|---|---|---|---|
| Degradation | $\gamma_{AR}$ | 0.1 | $h^{-1}$ |
| mRNA Production | $g_{AR}$ | 10 | mRNA $h^{-1}$ |
| Hill Coefficients | $n_{AR,\mu 200}$, $n_{AR,S}$, $n_{AR,Z}$ | 2, 2, 2 | Dimensionless |
| Hill Fold-Changes | $\lambda_{AR,\mu 200}$, $\lambda_{AR,S}$, $\lambda_{AR,Z}$ | 2, 0.5, 0.5 | Dimensionless |
| Hill Thresholds | $AR_{0,\mu 200}$, $AR_{0,S}$, $AR_{0,Z}$ | 90K, 90K, 90K | Molecules |

### 3 Simulations Details

In the single-cell formalism each isolated cell is described by the amount of the different proteins, TFs and miRs present in it as depicted in Figure 1A is always expressed in terms of number of molecules of each species. It is also possible that the cell might be exposed to external level of ligands associated with the Notch receptor like Delta $D_{ext}$ and Jagged $J_{ext}$ as well as Notch receptor $N_{ext}$, which simulates the scenario of cell-cell signaling between neighbour cells. The binning of the cells into different phenotypes is based on the level of miR200 (miR-200 < 5000 molecules: Mesenchymal; 5000<miR-200<15000: Hybrid Epithelial/Mesenchymal; miR-200>15000: Epithelial). These classifications holds true even at extreme values of the model parameters[Figure]. The single cell bifurcation plots were realised through the PyDSTool library of Python [6].

The multi-cell Tissue level lattice formalism was taken from the work of Boareto et al. [1]. It expands the single level formalism of Notch-EMT-AR into axis to a two-dimensional 60 by 60 lattice. The quantity of the molecular players are described by the same equations as described before for the single cell model. The tissue is modelled as a two-dimensional hexagonal lattice, with individual hexagons representing a cell. Each cell is bounded by six neighbouring cells and they can communicate with each other through Notch-Delta/Jagged mediated signling. At any given instance, the quantity of Notch receptor and its ligands which are free to bind to other cell's receptor and ligand is expressed as:

$$N_{ext} = \sum N_n \tag{29}$$

$$D_{ext} = \sum_n D_n + D_{sol} \tag{30}$$

$$J_{ext} = \sum J_n + J_{sol} \tag{31}$$

where $N_n$,$D_n$ and $J_n$ indicate the number of neighbour cells $n$ Notch receptors and ligands, and the total goes along each of the six connected neighbors. $D_{sol}$ and $J_{sol}$ indicate the amount of soluble Delta and Jagged that are present in the cellular matrix. This takes in account the paracrine activity that has been observed in several cancers [5] and the level of these soluble species remains constant throughout the simulation. The cells can also undergo EMT due to exposure to external EMT signal which directly activates Snail (see eq. 13). The equations governing the dynamics of the system were integrated numerically by an explicit Euler method having a time step of 0.1 hours, with a periodic boundary condition. The initial level of all the chemical species were set in a randomized order sampling homogeneously from the whole interval allowed for the bifurcation plots.

#### Supplementary Figures

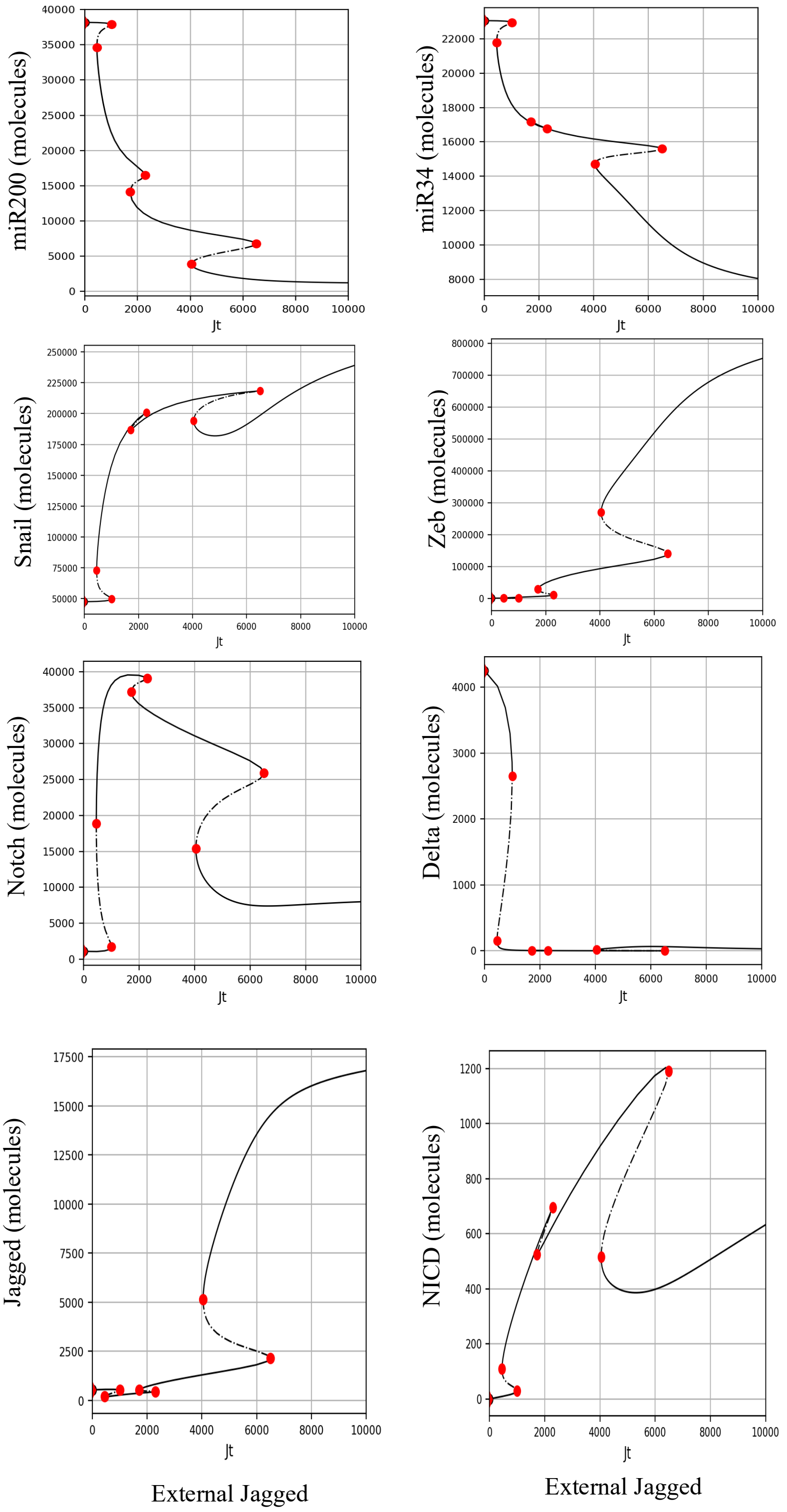


Figure 1: **Bifurcation plots of the chemical species present in the system in absence of AR as a function of external Jagged concentration.** Plots show the bifurcation of all the proteins and miRNAs present in the system corresponding to Figure 1C.

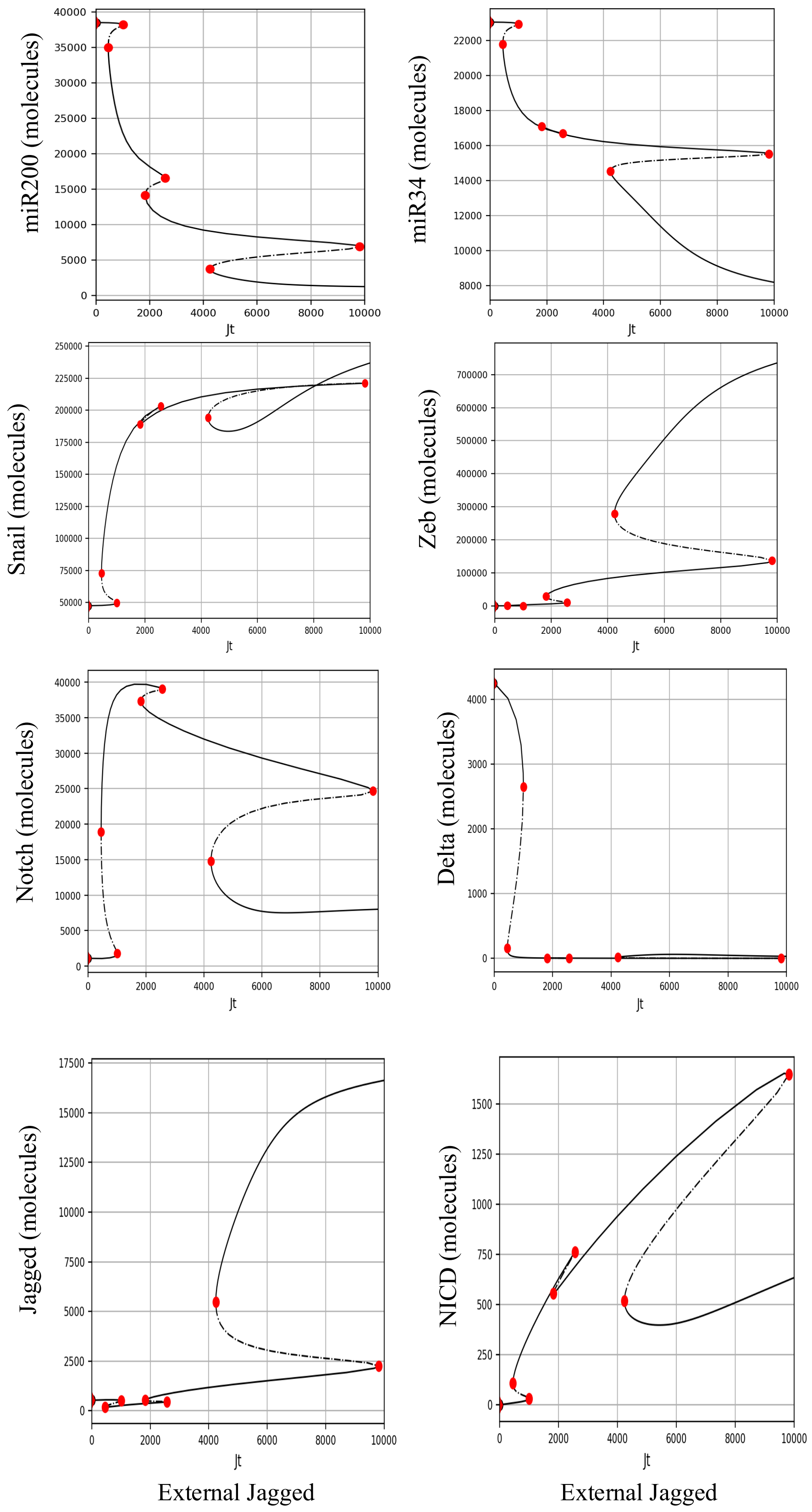


Figure 2: **Bifurcation plots of the chemical species present in the system in presence of AR as a function of external Jagged concentration.** Plots show the bifurcation of all the proteins and miRNAs present in the system corresponding to Figure 1D.

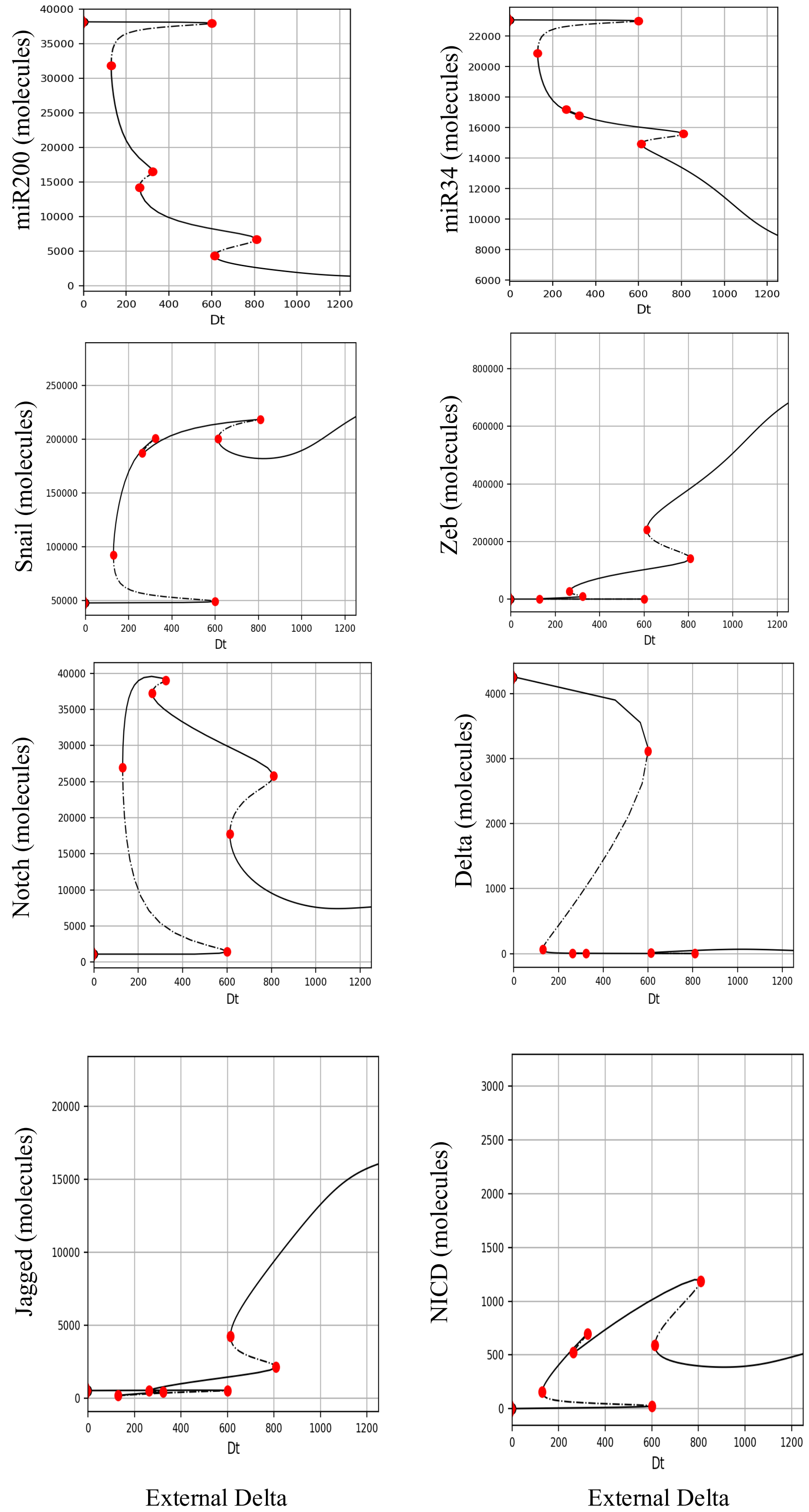


Figure 3: **Bifurcation plots of the chemical species present in the system in absence of AR as a function of external Delta concentration.** Plots show the bifurcation of all the proteins and miRNAs present in the system corresponding to Figure 1E.

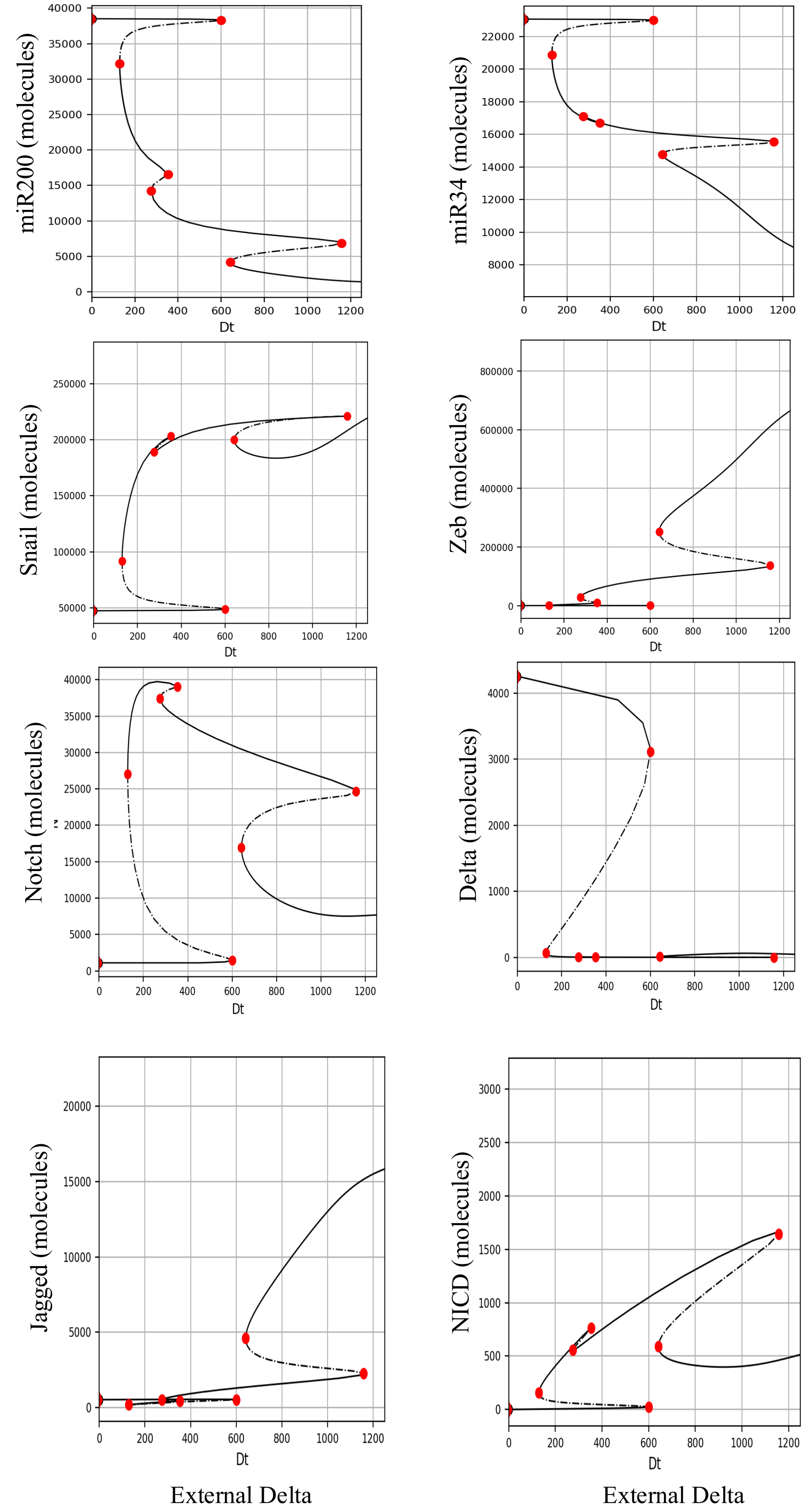


Figure 4: **Bifurcation plots of the chemical species present in the system in presence of AR as a function of external Delta concentration.** Plots show the bifurcation of all the proteins and miRNAs present in the system corresponding to Figure 1F.

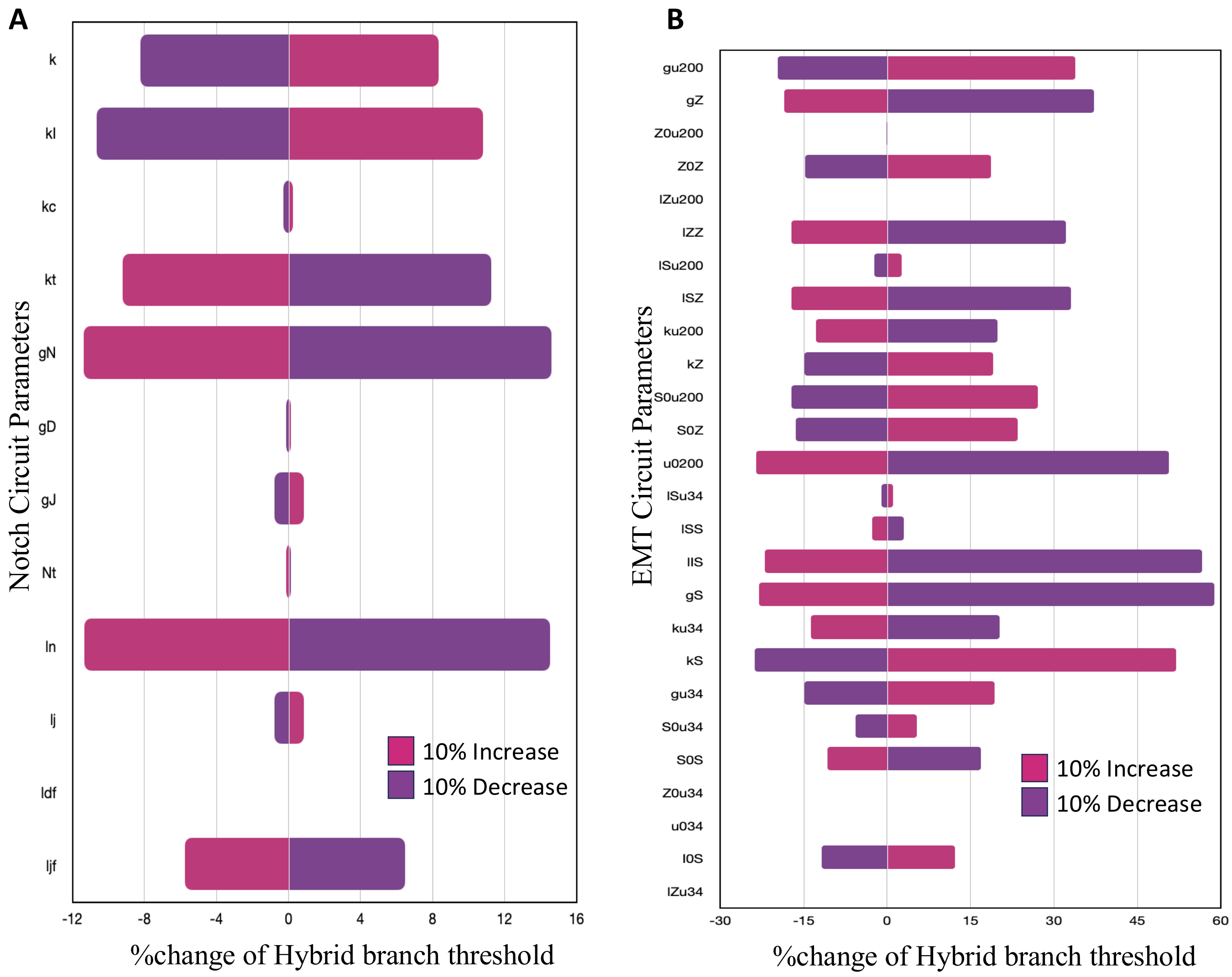


Figure 5: **Sensitivity Analysis of the single cell formulation under 10% perturbation of the system parameters.**(A)Variation in percentage of the minimum value of parameters required for the existence of a stable hybrid E/M branch in Figure 1D upon 10% variation of the variables belonging to the original Notch-Delta/Jagged circuit parameters of Jolly et al.[3]. (B) Sane as (A) but for the original EMT circuit parameters of Lu et al.[4].

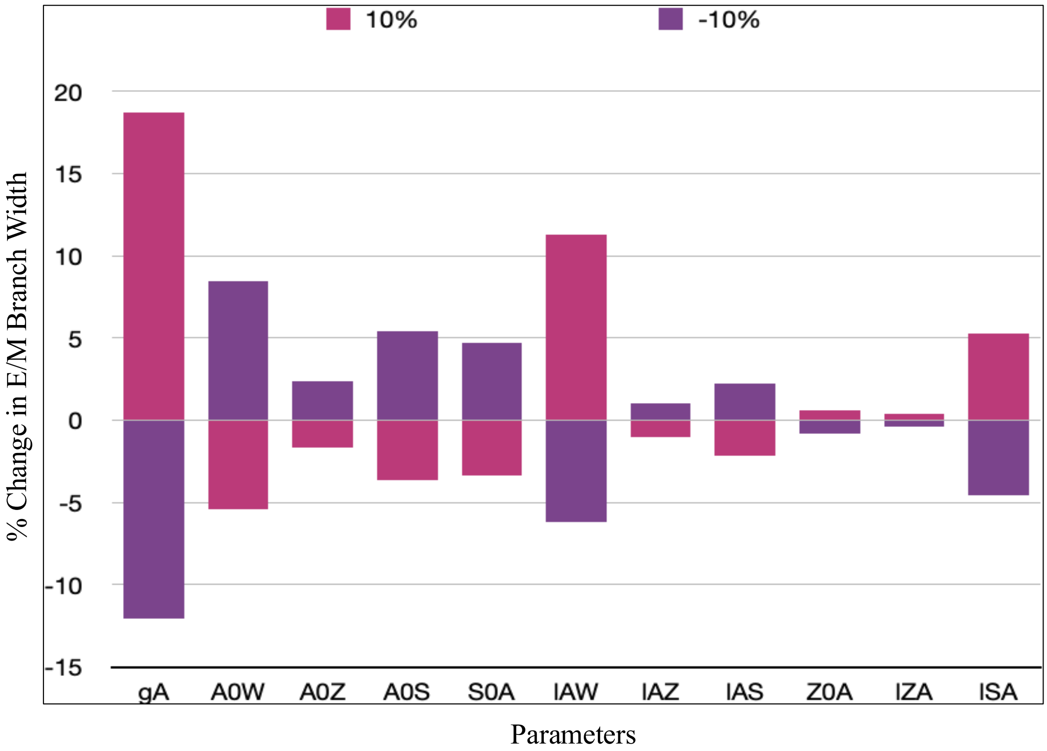


Figure 6: **Sensitivity Analysis of the single cell formulation under 10% perturbation of the system parameters associated with the interactions of Androgen Receptor [AR].** The percentage variation in the width of the hybrid E/M branch has been calculated on 10% variation of the parameters.

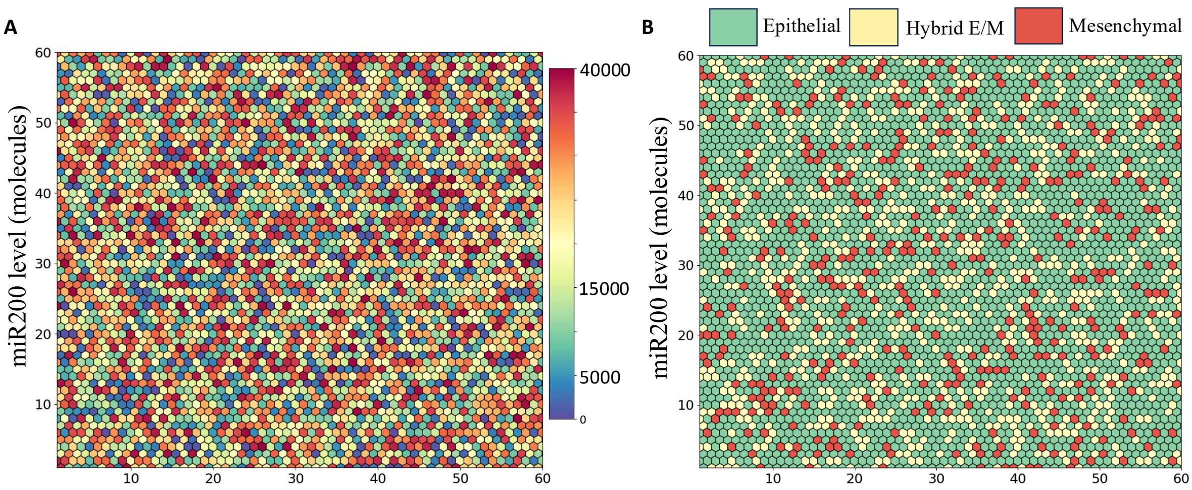


Figure 7: **Initial condition for the simulations presented in Figure 1 and Figure 2.** (A) Corresponding levels of all the chemical species were initialized randomly from an uniform distribution in the range of the maximum and minimum value of each species. The plot represents the level of miR200. (B) Phenotypic representation based on the level of miR200. E: miR200 $>$ 15000, E/M: 15000 $>$ miR200 $>$ 5000, M: miR200 $<$ 5000 molecules.

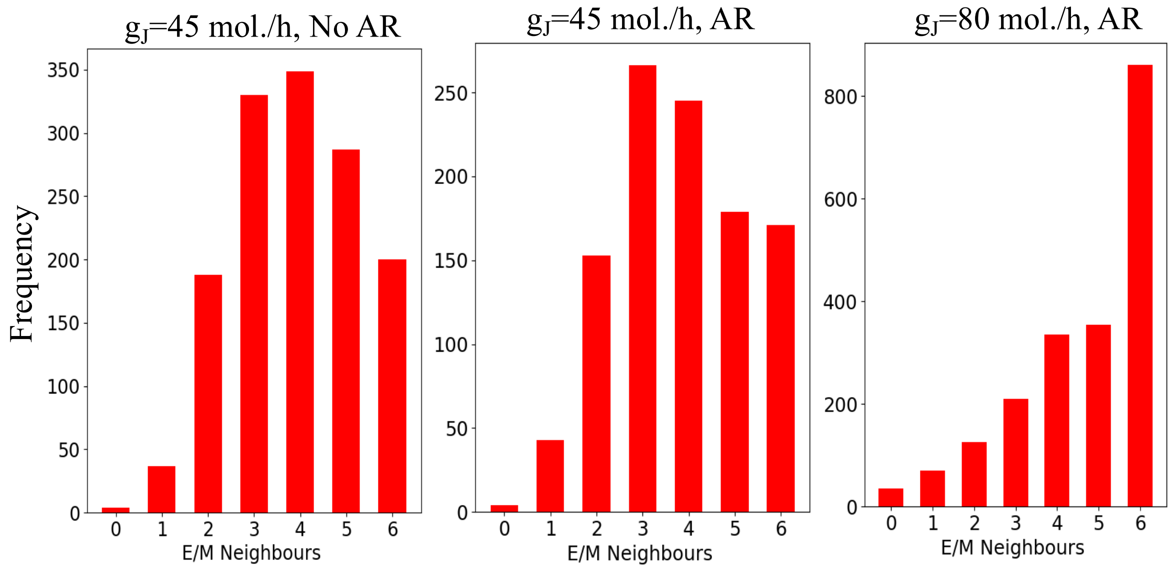


Figure 8: **Androgen Receptor increases the co-localization of the hybrid E/M phenotype.** Distribution of the hybrid E/M neighbours with respect to each hybrid E/M cell in the hexagonal lattice under diffeent production rates of Jagged corresponding to Figure 2 A, B, D.

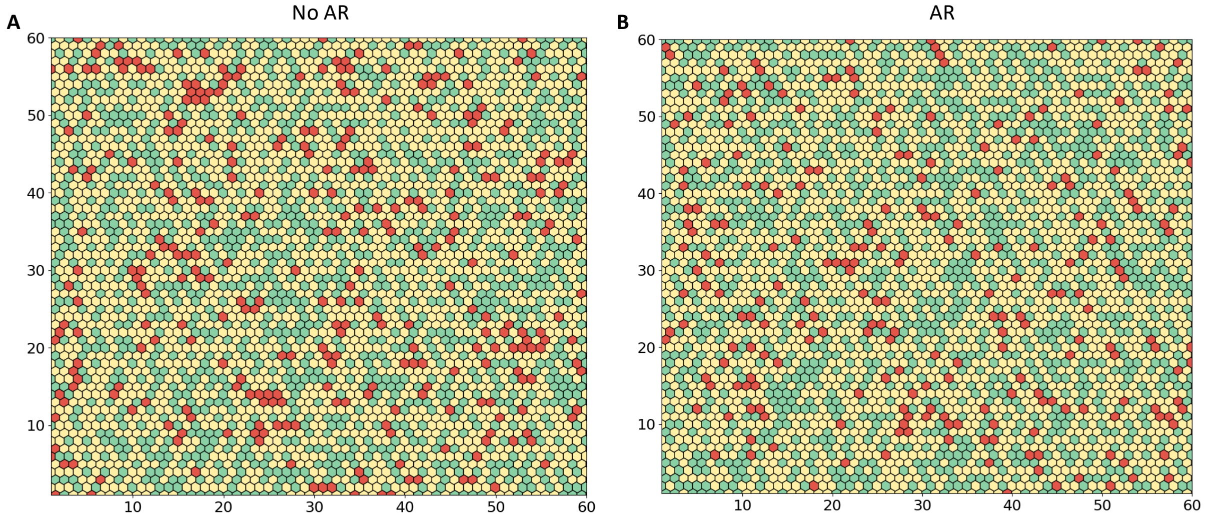


Figure 9: (A) Snapshot of the hexagonal lattice at 120 hours under $g_D = 85$ molecules/h and $g_J = 20$ molecules/h in the absence of AR. (B) Same as (A) in the presence of AR.

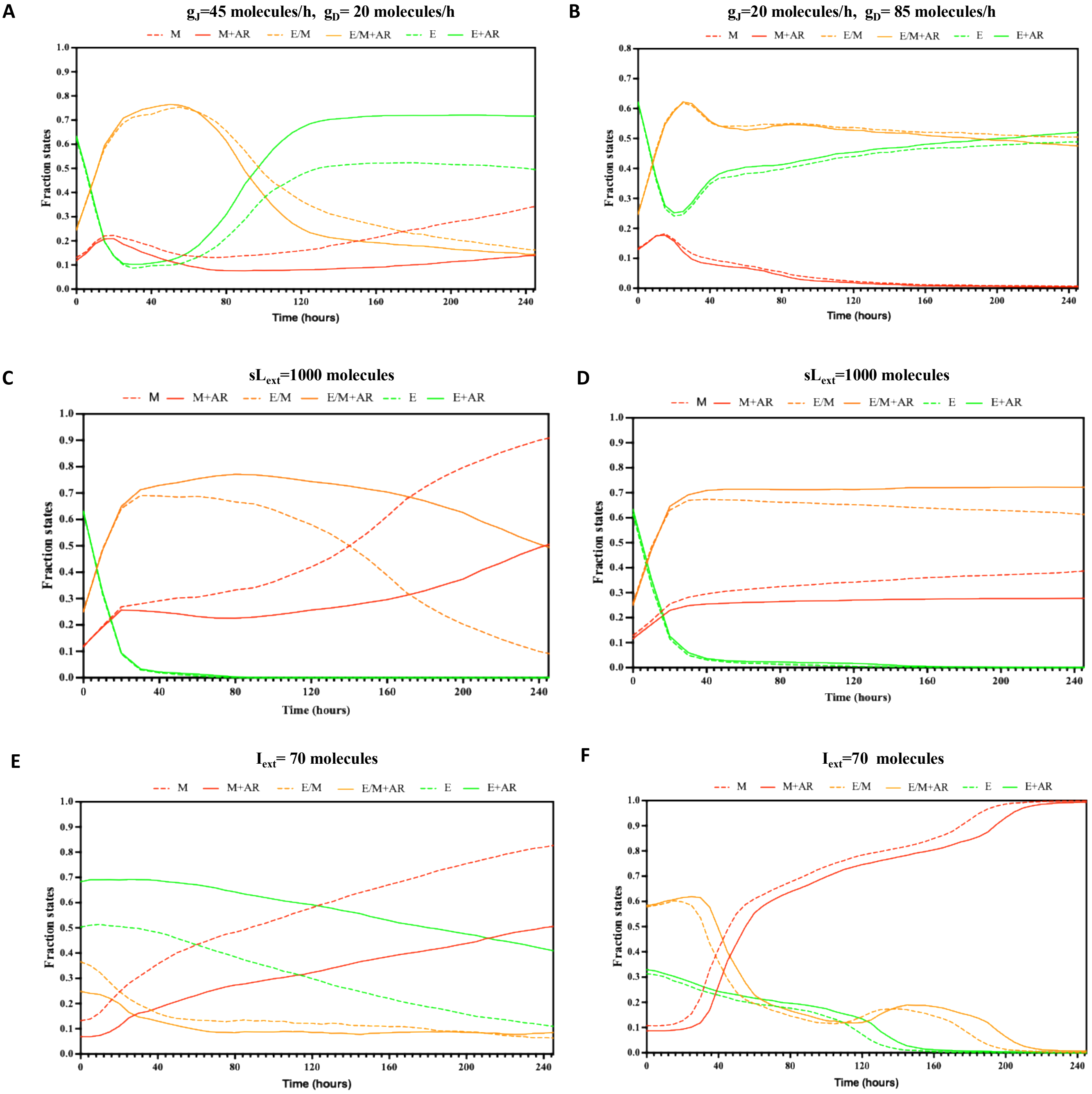


Figure 10: **Time evolution of the fraction of states under the influence of AR in the 60 by 60 lattice.** Plots in the left (A, C, F) are under a Jagged dominated signaling under $g_D = 20$ molecules/h and $g_J = 45$ molecules/h and on the right (B,D,F) under a Delta dominated signaling under $g_D = 85$ molecules/h and $g_J = 20$ molecules/h. In the plots A,B no external induction has been applied. Plot C,D were under the presence of external ligand concentration and plot E,F under the presence of external EMT signal $[I_{ext}]$. Plot A,B,C,D starts from the same initial randomized condition [Supplementary Figure 7]. Plot E starts from the configuration at the given level of Delta and Jagged without any induction in presence and absence of AR [Figure 2 A,B]. lot E starts from the configuration at the given level of Delta and Jagged without any induction in presence and absence of AR [Supplementary Figure 9 A,B]

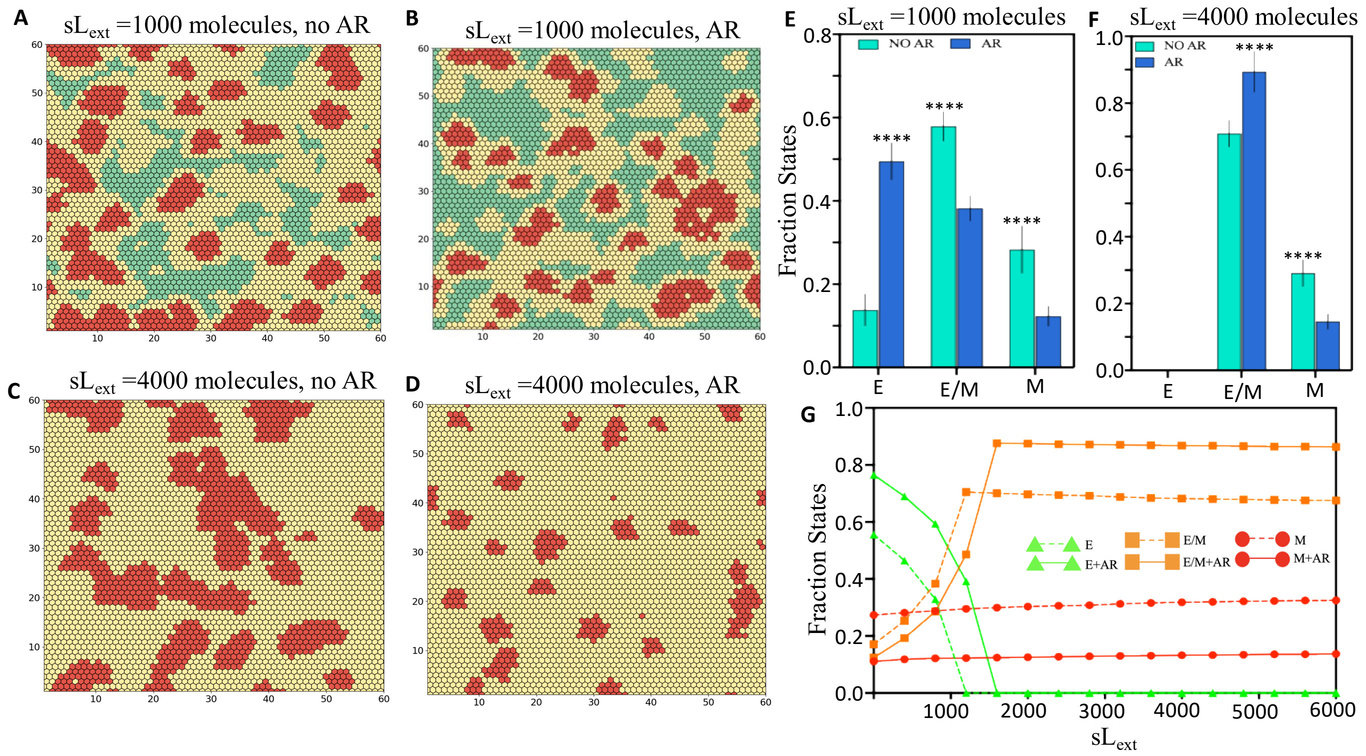


Figure 11: **Effect of AR on the tissue patterning of a Jagged dominated Notch signaling lattice under presence of soluble ligand.** (A) Snapshot of a two dimensional lattice without AR under $sL_{ext} = 1000$ molecules, (B) Same as (A) with AR, (C) Snapshot of a two dimensional lattice without AR at $sL_{ext} = 4000$ molecules, (D) Same as (C) with AR, (E) Average fraction of states for $sL_{ext} = 1000$ molecules, presence of AR increases fraction of cells in an epithelial state decreasing that for the hybrid E/M and mesenchymal state,(F)Average fraction of states for $sL_{ext} = 4000$ molecules, under high activation of the Notch signaling under this scenario all the cells are pushed towards a hybrid E/M and mesenchymal phenotype. Presence of AR increases the fraction of state in hybrid E/M phenotype decrease that for the mesenchymal state (G) Fraction of E, hybrid E/M and M cells as a function of the concentration of external ligand $[sL_{ext}]$ in a two dimensional lattice in the presence and absence of AR. In the lattice, green represents epithelial state, yellow stands for hybrid state and red stands for mesenchymal state. The production rate of Jagged was fixed at $g_J = 45$ molecules/h and Delta was fixed at $g_D = 20$ molecules/h for all the simulations. Each simulations were started from the state obtained in Figure 2 A,B in the presence and absence of AR respectively. For each simulation the snapshots and the fraction of states were calculated at 120 hours. For statistical analysis number of replicates were 10. The significance figures are given in the methodology section.

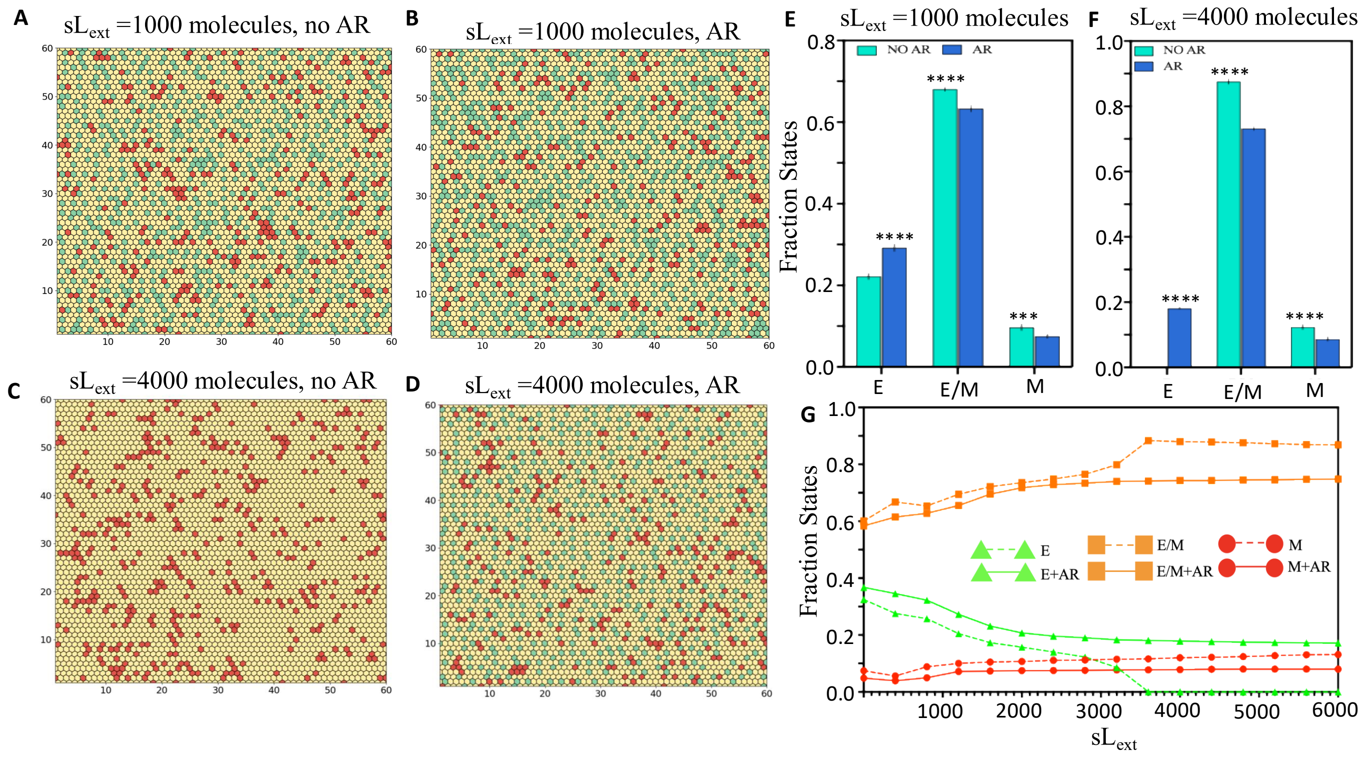


Figure 12: **Effect of AR on the tissue patterning of a Delta dominated Notch signaling lattice under presence of soluble ligand.** (A) Snapshot of a two dimensional lattice without AR under $sL_{ext} = 1000$ molecules, (B) Same as (A) with AR, (C) Snapshot of a two dimensional lattice without AR at $sL_{ext} = 4000$ molecules, (D) Same as (C) with AR, (E) Average fraction of states for $sL_{ext} = 1000$ molecules, presence of AR increases fraction of cells in an epithelial state decreasing that for the hybrid E/M and mesenchymal state,(F) Average fraction of states for $sL_{ext} = 4000$ molecules, under high activation of the Notch signaling under this scenario all the cells tend to attain a hybrid E/M or mesenchymal phenotype. But presence of provides stability to all the states, hence maintaining a constant fractional distribution, (G) Fraction of E, hybrid E/M and M cells as a function of the concentration of external ligand $[sL_{ext}]$ in a two dimensional lattice in the presence and absence of AR. In the lattice, green represents epithelial state, yellow stands for hybrid state and red stands for mesenchymal state. The production rate of Jagged was fixed at $g_J = 20$ molecules/h and Delta was fixed at $g_D = 85$ molecules/h for all the simulations. Each simulations were started from the state obtained in Figure 3 A,B in the presence and absence of AR respectively. For each simulation the snapshots and the fraction of states were calculated at 120 hours. For statistical analysis number of replicates were 10. The significance figures are given in the methodology section.

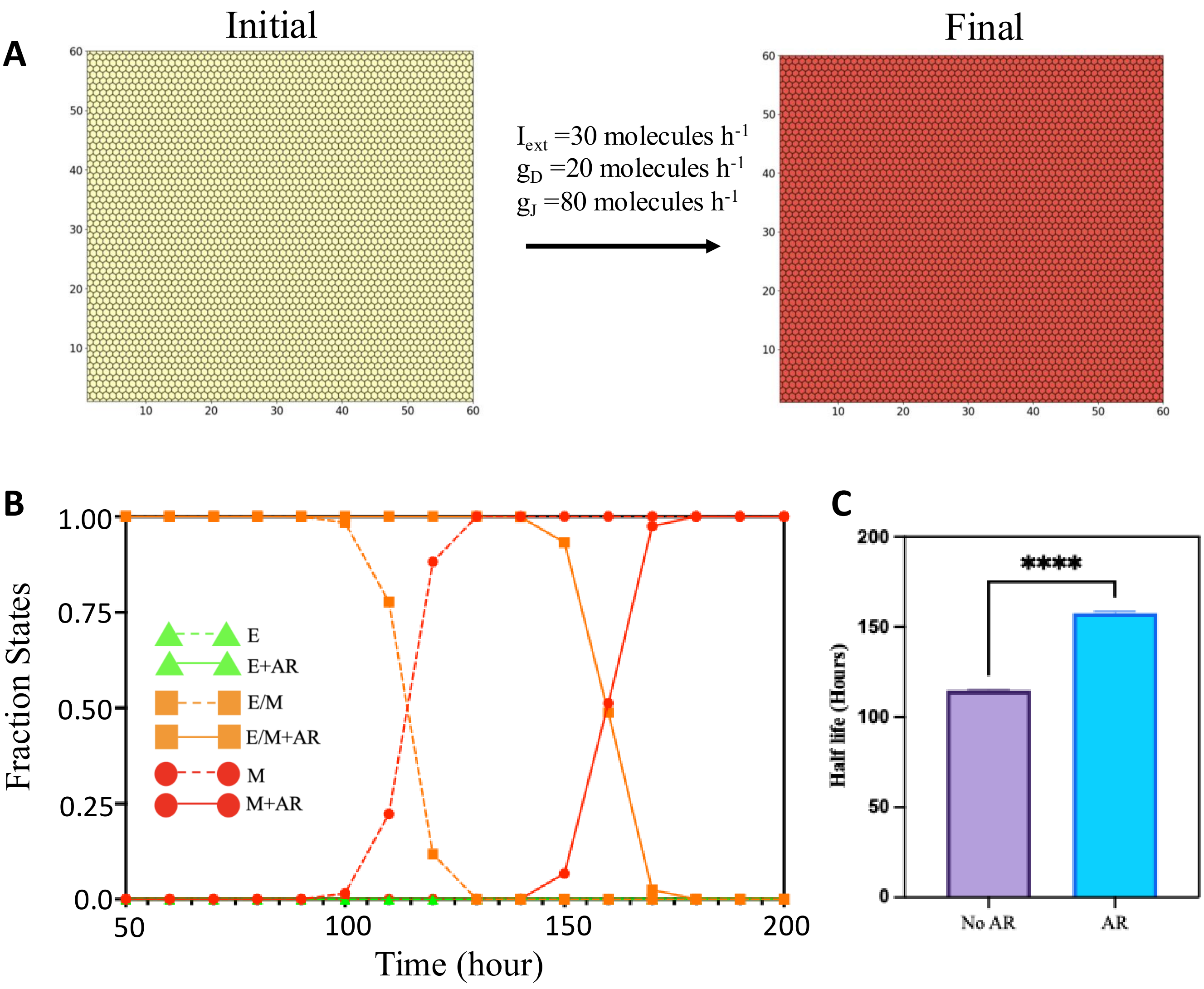


Figure 13: **Presence of AR provides temporal stability to the hybrid E/M phenotype under Notch-Jagged signaling.** (A) The cell lattice was initialized from sampling out randomly from the ranges of the chemical species that allows the existence of a hybrid E/M state. It was then exposed to an EMT inducing signal $I_{ext}$ through Snail, which pushed the cells towards a mesenchymal state under a Jagged mediated Notch signaling condition. (B) The fraction of state plot of the different phenotypes with respect to time. The transition points where the ratio between the hybrid E/M state and mesenchymal state is 1:1 is considered as the half life of the hybrid E/M state. (C) Bar plot representing the half life of the hybrid E/M state. The presence of AR significantly enhances the half life of the hybrid E/M state. The number of replicates for statistical analysis was 10. The significance figures are given in the methodology section.

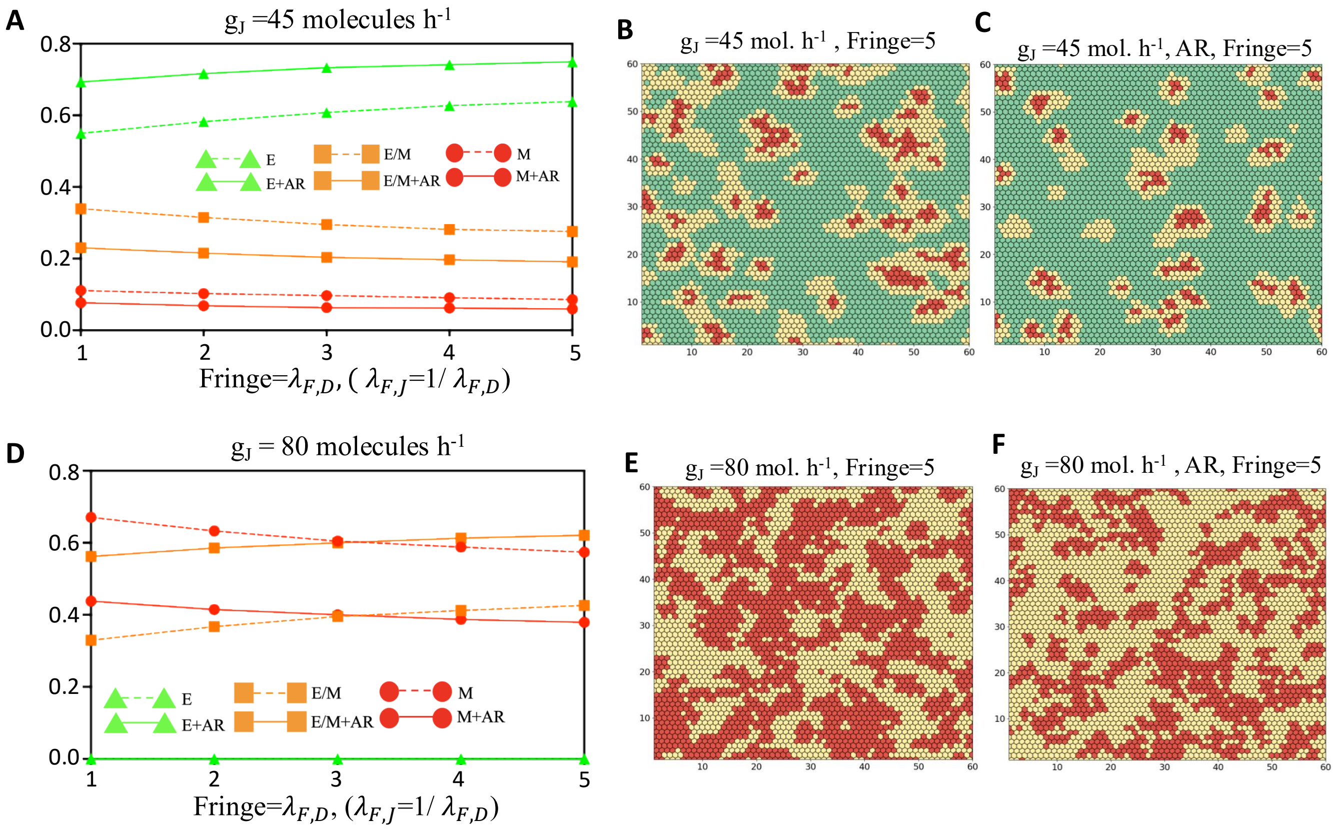


Figure 14: **Effect of Fringe on the tissue patterning and phenotypic distribution in the tissue lattice in presence and absence of AR under Jagged dominated signaling.** (A) Fraction of states distribution of the three phenotypes epithelial (E), hybrid E/M (E,M) and mesenchymal (M) in the presence and absence of AR with respect to varying strength of Fringe modulation on Notch-Delta and Notch-Jagged signaling in a 60 by 60 lattice after 120 hours with production rate of Jagged fixed at $g_J$ = 45 molecules/h. (B) Snapshot of the tissue lattice at 120 hours under the Fringe strength of 5 in the absence of AR. The cluster of hybrid E/M and mesenchymal cells decreases as compared to Figure 2A. (C) Same as (B) in the presence of AR. (D) Same as (A) under the production rate of Jagged set to $g_J$ = 80 molecules/h. (E) Same as (B) under the production rate of Jagged set to $g_J$ = 80 molecules/h. The Fringe modulation increases the number of cells in a hybrid E/M phenotype [compare with Figure 2C] by decreasing the strength of the Notch signaling through the decrease in affinity of Notch towards Jagged and increase in affinity towards Delta. (F) Same as (E) under the presence of AR. In all the above scenarios the production rate of Delta was fixed at $g_D$ = 20 molecules/h.

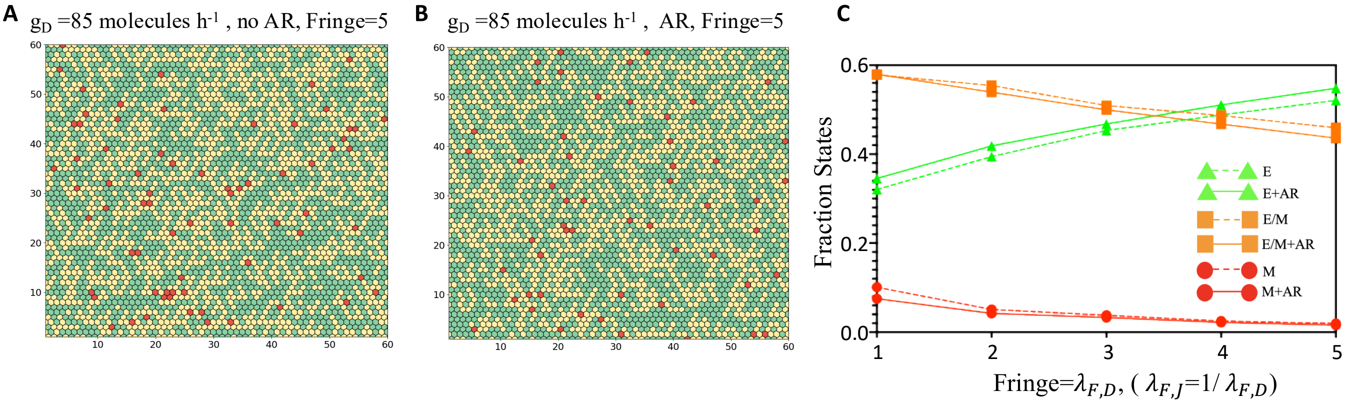


Figure 15: **Effect of Fringe on the tissue patterning and phenotypic distribution in the tissue lattice in presence and absence of AR under Delta dominated signaling.** (A) Snapshot of the tissue lattice at 120 hours in the absence of AR under the Fringe modulation of 5. Fringe increases the fraction of cells in the epithelial phenotype [compare with Figure 3C]. (B) Same as (A) in the presence of AR. Under both the presence and absence of AR, Fringe modulation of the Notch-Delta and Notch-Jagged interaction can increase the fraction of cells in epithelial state by reducing the activation of the Notch signaling. (C) Fraction of states distribution of the three phenotypes epithelial (E), hybrid E/M (E,M) and mesenchymal (M) in the presence and absence of AR with respect to varying strength of Fringe modulation on Notch-Delta and Notch-Jagged signaling in a 60 by 60 lattice after 120 hours. In this case the production rate of Delta was fixed at $g_D = 85$ molecules/h and that of Jagged was fixed at $g_J = 20$ molecules/h.

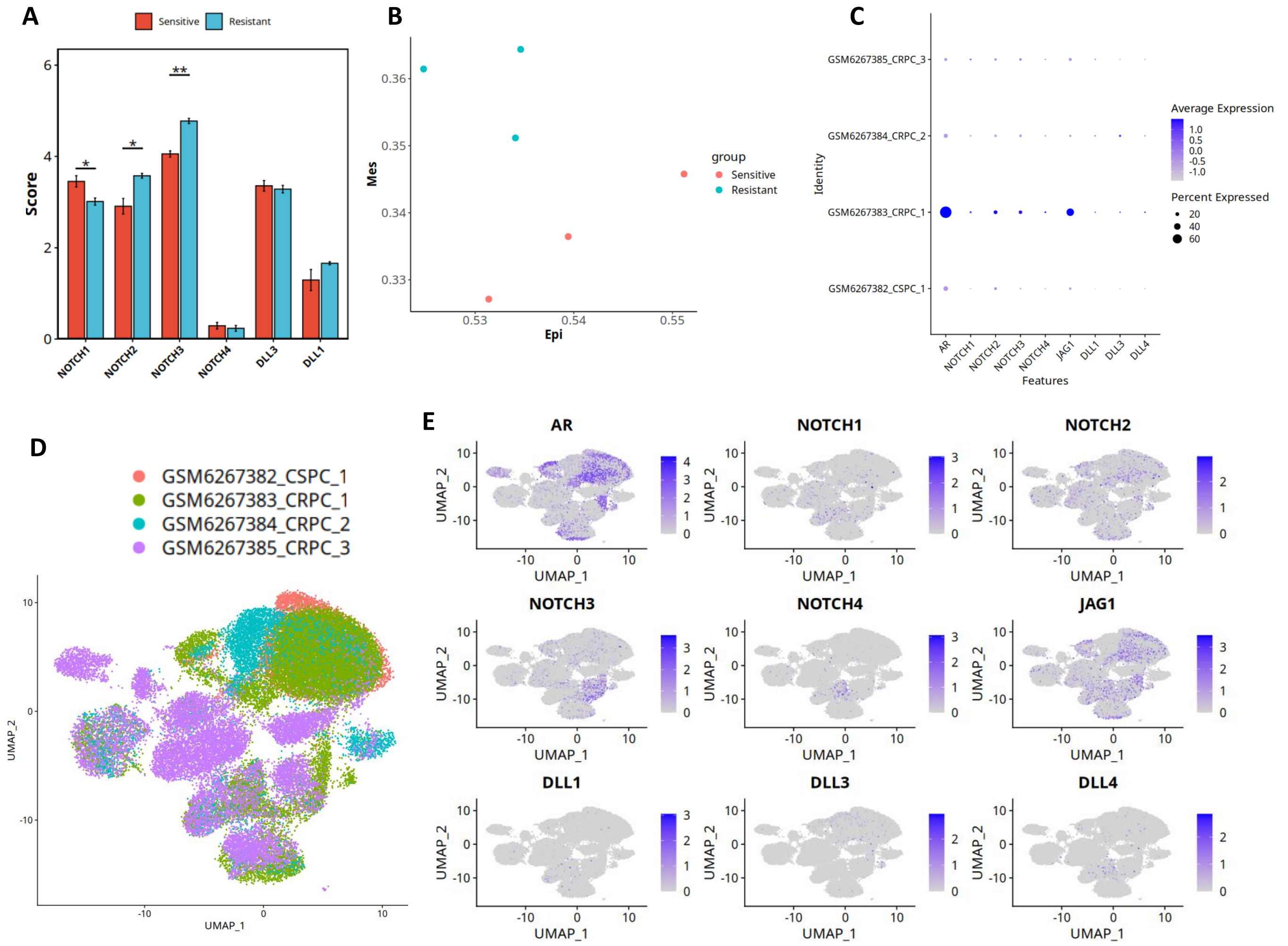


Figure 16: **Gene expression patterns in in vitro and patient data for androgen sensitive vs resistant prostate cancer.** (A) Boxplots showing expression of NOTCH and Delta ligands expression for sensitive and resistant cell lines from GSE123379 [7]. (B) 2D scatter plot depicting the Epithelial and mesenchymal status of the sensitive and resistant samples. (C). Dot plot showing the average expression of AR, JAG1, NOTCH and Delta ligands across four single cell RNAseq patient data. (D). Dim plot to visualize the single-cell RNA-seq data after UMAP reduction. The cells are coloured by sample type to visualize cell clustering based on resistance to androgen. (E). Featureplot to visualize distribution of gene expression of AR, JAG1, NOTCH and Delta ligands on dimensionally reduced clusters.

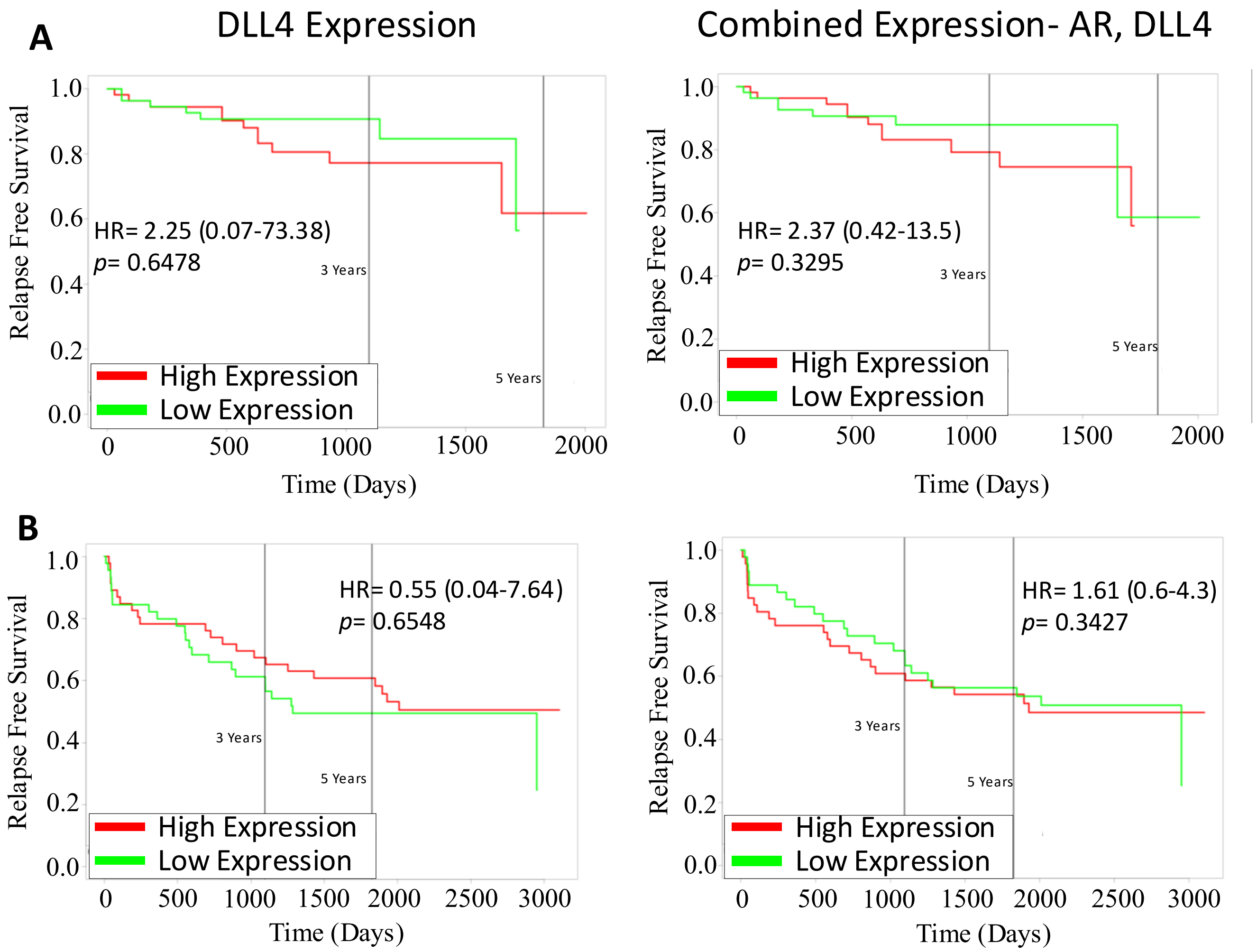


Figure 17: **Survival analysis for co-expression of AR and DLL4 in several PCa patient cohorts.** (A) Kaplan–Meier curves for overall survival (OS) for DLL4 high vs low expression (left) and both AR and DLL4 combined high vs low expression (right) in prostate cancer patient data from GSE70768 . Reported p-values are based on a log-rank test a indicating significant difference in survival. Same as (A) but for (B), Relapse Free Survival in prostate cancer patient data from GSE70769.
